# Tryptophan degradation by intestinal Bacteroides induces anti-tumor immunity and limits melanoma growth

**DOI:** 10.64898/2026.05.06.723300

**Authors:** Ximena Diaz Olea, Kristin Beede, Gabriel Pereira, David Scott, Christopher Petucci, Eric Martens, Dmitri Rodionov, Aagam Shah, Miguel P. Martinez, Hyungsoo Kim, Ashok Kumar Sharma, Anthony Martin, Tongwu Zhang, Mark B. Faries, Omid Hamid, Suzanne Devkota, Andrei Osterman, Simon Knott, Emile E. Voest, Nadim J. Ajami, Jennifer Wargo, Amanda E. Ramer-Tait, Ze’ev A Ronai

## Abstract

Defining mechanisms used by gut microbiota to control anti-tumor immunity may offer novel therapeutic modalities. Here, we demonstrate that *Bacteroides rodentium* and closely related *Bacteroides uniformis* species induce anti-tumor immunity and limit melanoma development when colonized in either germ-free (GF) mice, mice with a complex microbiome, or WT mice. Enhanced CD8^+^ T cell infiltration seen in tumors of mice harboring *B. rodentium* coincided with increased expression of immune-stimulating pathways and activation of bone marrow-derived dendritic cells that were co-cultured with the *B. rodentium* secretome. Metabolomic analyses of cecal samples from GF mice colonized with Altered Shedlar Flora (ASF) plus *B. rodentium* revealed lower tryptophan levels than in ASF-colonized controls, and WT mice fed a tryptophan-deficient diet exhibited inhibition of melanoma development. *In silico* genomic reconstruction of metabolic pathways revealed that both *B. rodentium* and *B. uniformis* harbor tryptophanase A (*TnaA*) and aromatic amino transferase (*ArAT*) genes, both of which function in tryptophan degradation. Administration of a *B. uniformis* harboring *TnaA* mutant failed to inhibit melanoma growth in gnotobiotic mice. Notably, administration of indoles, but not kynurenines, also effectively inhibited melanoma development, increasing immune cell infiltration into the tumors. Correspondingly, levels of bacterially encoded tryptophan-degrading enzymes were higher in cohorts of melanoma patients responding to immune checkpoint blockade. These findings highlight a novel mechanism of anti-tumor immunity and tumor growth inhibition dependent on the tryptophan degradation products, indoles, produced by intestinal *Bacteroides* species.

## Introduction

Melanoma, which exhibits high metastatic propensity and resistance to therapy, is one of the most aggressive skin cancers. Although combining immunotherapy with targeted therapies holds great promise against melanoma^1,2^, alternatives that overcome immune evasion and enable immune system recognition of immune-cold tumors remain unmet needs. Among options to address these needs are strategies to selectively alter the gut microbiota, which play an important role in controlling melanoma development and enhancing responses to chemotherapy, radiation, or immunotherapies^3–7^. In mouse models, strategies to enrich for specific gut bacteria, such as *Bacteroides* in melanoma^4^ or *Faecalibaculum* and *Bacteroides* in colorectal cancer (CRC)^8^, reportedly attenuate tumor growth, although underlying mechanisms remain unknown.

Efforts to define mechanisms regulating anti-tumor immunity led us to investigate RNF5, a membrane-anchored E3 ubiquitin ligase that functions in Endoplasmic Reticulum Associated Protein Degradation (ERAD)^9^. RNF5 serves to clear misfolded proteins such as CFTR, SLC1A5 and S100A8, which are implicated in pathogenesis of multiple chronic diseases, including inflammatory bowel diseases and cancer^10–12^. Notably, we found that *Rnf5* knock out (KO) mice exhibit limited tumor growth due to enhanced anti-tumor immunity driven by changes in the gut microbiota^4^. Administration of 11 bacterial strains enriched in *Rnf5* KO mice was sufficient to induce anti-tumor immunity and inhibit melanoma growth in GF mice^4^.

Mechanisms underlying regulation of anti-tumor immunity are under intense investigation. Growing evidence indicates that metabolic activities play a key role in anti-tumor immunity, with particular attention to the amino acid tryptophan, which may antagonize anti-tumor immunity^13–15^. Notably, analysis of this activity has largely focused on cellular changes in both cancer and immune cells, although the possible role of a more systemic cause remains plausible. In humans and mice, tryptophan must be obtained exclusively through diet. Tryptophan is also a precursor to diverse metabolites produced by numerous intestinal microorganisms^14^ that have been implicated in intestinal immune tolerance and gut microbiome homeostasis^13–15^. Numerous studies report that certain bacterial species can also activate subsets of immune cells, either directly or indirectly, resulting in varying degrees of anti-tumor immune activity^5,16–18^. The amino acid tryptophan is implicated in these activities and has been shown to negatively regulate immune system factors and block anti-tumor immunity, largely by regulating activity of the aryl hydrocarbon receptor (AhR) and indoleamine 2, 3-dioxygenase 1 (IDO1)^19–21^.

Here, we aimed to define specific gut bacterial species and related metabolites required for anti-tumor immunity and melanoma growth inhibition using mouse models with varying microbiome complexity^22,23^. Our analyses identified *B. rodentium,* one of the 11 bacterial strains we previously reported to be enriched in the gut microbiome of *Rnf5* KO mice, and a closely related species found in humans, *B. uniformis,* as capable of enhancing anti-tumor immunity and limiting melanoma growth. Moreover, we show that in *B. uniformis,* deletion of *tryptophanase A* (*TnaA*), which encodes an enzyme catalyzing tryptophan degradation, blocked induction of anti-tumor immunity and its ability to inhibit melanoma growth. Collectively, these studies establish that tryptophan degradation by the gut microbiota is an important process in controlling anti-tumor immunity and tumor growth.

## Results

### B. rodentium induces anti-tumor immunity and inhibits melanoma growth

Our earlier studies identified 11 bacterial species that enhanced anti-tumor immunity and inhibit melanoma development when provided to germ-free (GF) mice harboring the Altered Schaedler Flora (ASF)^4^. Analyses identified three bacterial strains - *Bacteroides rodentium, Phocaicola sartonii*, and *Phocaicola massillennsis* - that were significantly enriched at the end of this study (tumor collection point) (Figure 1A). Of those, colonization of *Bacteroides rodentium* (*B. rodentium*) with ASF promoted immune development in GF recipients, inhibiting melanoma growth as evidenced by decreased tumor size compared to ASF controls (Figure 1B, Supplemental Figure 1C). However, administering *P. sartorili* and *P. massiliensis* abrogated anti-tumor effects mediated by *B. rodentium* (Supplemental Figures 1A,1B). Importantly, colonizing MC608-F-a1 mice harboring a more complex microbiota deficient in *Bacteroides*^22–24^ with *B. rodentium*, was also sufficient to inhibit melanoma growth (Figure 1C, Supplemental Figure 1D). Notably, the presence of the *Bacteroides* genus was associated with better overall survival in the TCGA solid-tumor cohort (Supplemental Figure 1E).

**Figure 1:**
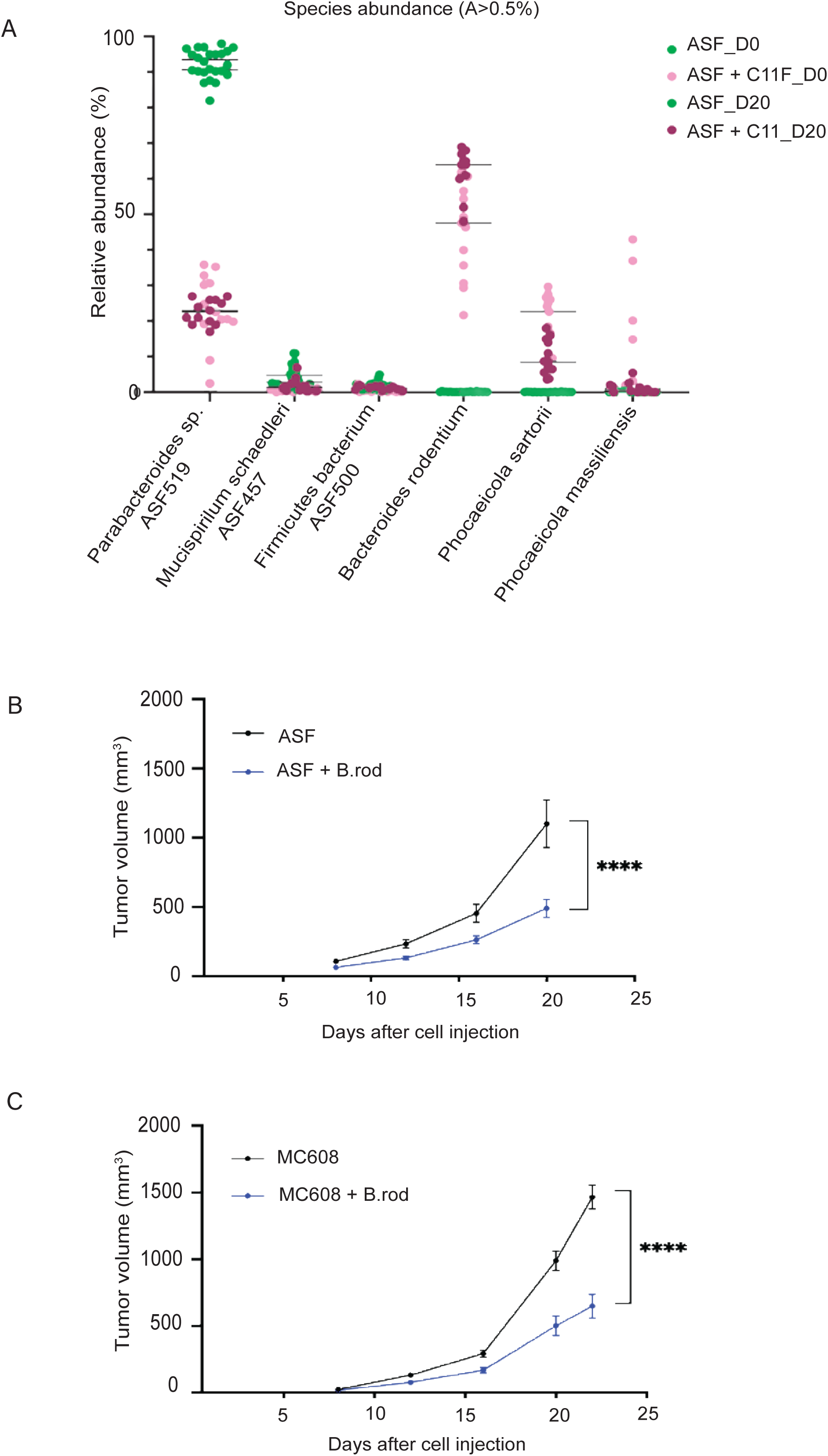
*B. rodentium* limited tumor growth. **(A)** Average relative abundance (A%) >0.5% of bacterial species in the gut microbiomes of germ-free (GF) mice administered the Altered Schaedler Flora (ASF) or ASF plus a cocktail of 11 bacterial species on days 0 and 20 days after subcutaneous injection of YUMM 1.5 cells. **(B)** Tumor growth in GF mice colonized with either ASF or ASF plus *B. rodentium* via oral gavage 14 days prior to YUMM1.5 tumor cell injection (n = 15 mice/treatment; data represent two experiments). **(C)** Tumor growth in GF mice colonized with either microbiome MC608-F-a1 or MC608-F-a1 plus *B. rodentium* 14 days prior to YUMM1.5 tumor cell injection (n=18 mice/treatment; data represent two experiments). Data were analyzed by unpaired t-test. ****P < 0.0001 using two-way ANOVA.

### B.rodentium colonized mice show immune cell infiltration of tumors

To assess gene expression changes associated with anti-tumor immunity, we performed RNAseq of tumor tissues collected from GF mice colonized with ASF in the presence or absence of *B. rodentium*. Notably, relative to controls receiving only ASF, mice harboring *B. rodentium* exhibited enhanced expression of genes associated with immune activation (Figure 2A), including CD86 and ICAM (implicated in T cell receptor signaling), CCR1, TLR1, and TLR2 (implicated in cytokine signaling), and several cytokines, including TNFα and IFNγ (Figure 2A and Supplemental Figure 2A-B). RT-qPCR analysis confirmed upregulation of these genes in tumors obtained from mice colonized with *B. rodentium* versus controls (Supplemental Figure 2A-B). Further analysis showed enhanced CD8^+^ T cell infiltration of tumors obtained from mice colonized with *B. rodentium* compared to ASF controls (Figure 2B). To test whether *B. rodentium* alters the gut epithelial barrier, we examined the architecture of the small intestine in MC608-F-a1 mice colonized with *B. rodentium.* We noted shorter villi and increased number of crypts in mice colonized with *B. rodentium* compared to MC608-F-a1 controls (Figure 2C), similar to outcomes seen in mice that were previously reported to exhibit enhanced anti-tumor immunity^4^.

**Figure 2:**
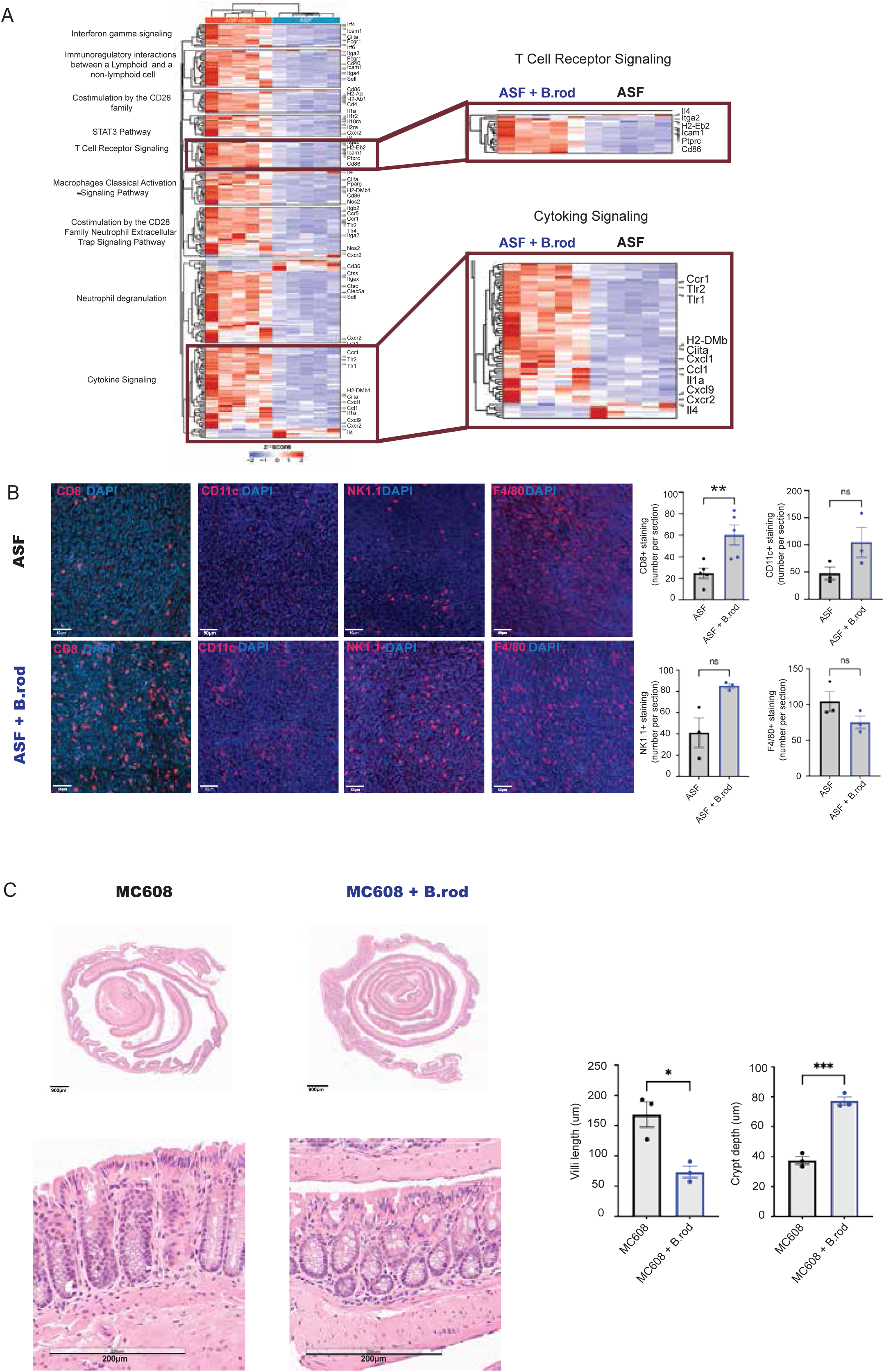
Upregulated immune signaling in mice colonized with *B. rodentium*. Gene expression changes were assessed by RNAseq (on day 22) in tumor samples from GF mice colonized with ASF or ASF plus *B. rodentium* (n = 6 mice/treatment; two independent experiments). **(A)** Bioinformatic analysis following RNAseq was performed on tumors from GF mice colonized with ASF or ASF plus *B. rodentium*. Heatmap shows differentially expressed pathways. **(B)** Staining with anti-mouse CD8, CD11c, NK1.1, and F4/80 antibodies of tumor sections described above. CD8^+^, CD11c^+^, NK1.1^+^, and F4/80^+^ positive cells were quantified (n=3 mice/treatment in two experiments). For image analysis, 3 images were randomly selected from different areas of each tumor, in which the number of positive cells per section were counted. Presented is the average number of these three sections. **(C)** Staining with H&E of Swiss roll sections from small intestines obtained from mice colonized with MC608-Fa1 or MC608-Fa1 plus *B. rodentium.* (scale bar 200 µm). Villi lengths and crypt depth were measured (n=3 mice/treatment in two experiments). Data were analyzed by unpaired t-test. *P < 0.05, **P < 0.005, ***P < 0.001 using two-tailed t-test or Mann-Whitney U test.

To assess how *B. rodentium* impacts immune cell function, we collected the secretome from *in vitro* cultures of *B. rodentium* and incubated it with bone marrow-derived dendritic cells (BMDCs) from naïve mice. BMDCs exposed to the *B. rodentium* secretome showed increased expression of markers of dendritic cell (DC) activation such as CD80, CD40 and MHC I relative to untreated BMDCs (Supplemental Figure 2C). These observations suggest that *B. rodentium* can activate immune cells that function in antitumor immunity.

### Tryptophan depletion inhibits melanoma growth

To assess mechanisms underlying anti-tumor activity by *B. rodentium*, we performed metabolomic analysis of cecal samples from mice colonized with ASF plus *B. rodentium* compared to ASF only controls (Supplemental Table 3). Strikingly, we observed significantly reduced levels of tryptophan in cecal samples of mice colonized with *B. rodentium* plus ASF compared with ASF only (Figure 3A). Mice harboring *B. rodentium* also showed decreased cecal levels of tyrosine and isoleucine and increased levels of the host-derived metabolites dimethylarginine, hydroxyproline and taurine (Figure 3A). Independent metabolomic analyses confirmed increased metabolites associated with tryptophan metabolism (Supplemental Figure 3A). Consistent with these observations, changes in the level of tryptophan metabolites produced by the bacterial enzymes were among the more pronounced (Supplemental Figure 3B). Correspondingly, elevated levels of tryptophan degradation products, such as quinolinic acid and kynurenic acid, which are derived upon tryptophan degredation by the host were also identified (Supplemental Figure 3C).

**Figure 3:**
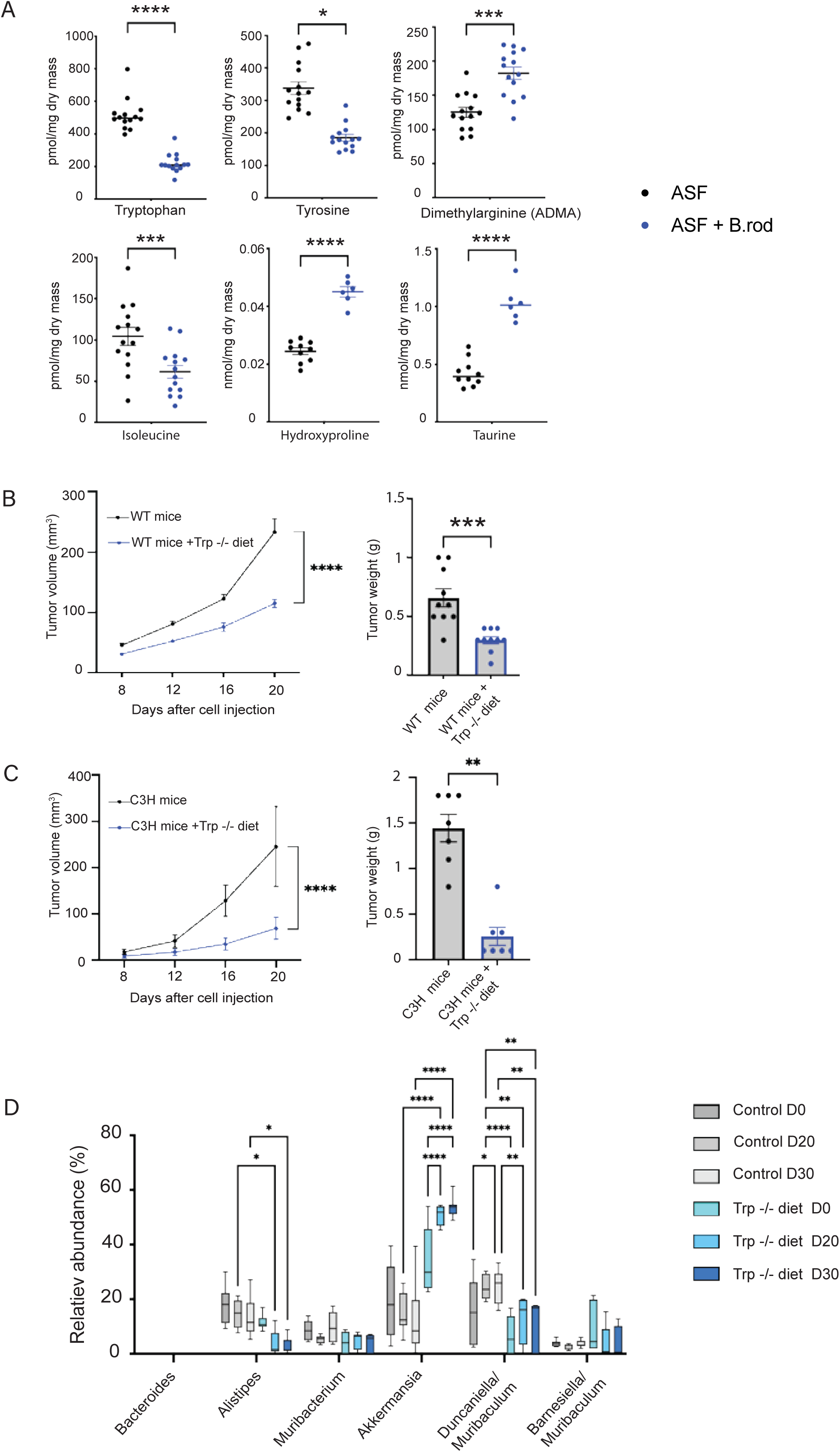
Lower levels of tryptophan were identified in metabolomic analysis of cecal samples from gnotobiotic mice. **(A)** Quantification of indicated metabolites in cecal samples from GF mice colonized with ASF or ASF plus *B. rodentium* (n = 6 mice/treatment). **(B)** Growth and weight of tumors in conventional C57BL/6J mice or **(C)** C3H mice. Provided with the indicated diets (control or tryptophan-deficient) three days prior to YUMM1.5 tumor cell injection (day 0; n = 10 mice/treatment). **(D)** Taxonomic phenotype profile of bacterial strains was obtained at the indicated time points from fecal samples. Analyses were based on 16S rRNA bacterial gene sequencing followed by bioinformatic analyses. Data were analyzed by unpaired t-test. *P < 0.05 **P < 0.005, ***P < 0.001, ****P < 0.0001 by two-tailed t-test or two-way ANOVA.

Supplementing the diet of conventional C57BL/6J mice with either kynurenic acid or taurine, both previously shown to affect anti-tumor immunity in independent studies^25,26^, did not alter tumor development (Supplemental Figures 3D and 3E). Further, conventional C57BL/6J or C3H mice fed a tryptophan-deficient diet starting three days prior to tumor cell injection showed significant inhibition of melanoma development compared to mice fed a diet containing tryptophan (Figure 3B and 3C). Notably, these mice also showed marked changes in gut bacterial composition (Supplemental Figure 3F), the most notable being a significant enrichment in *Akkermansia municiphila* (Figure 3D), a tryptophan prototroph strain previously associated with positive responses in patients to immune checkpoint therapies^27^. At the same time, the relative abundance of *B. rodentium* was unaltered (Supplemental Figure 3G), suggesting that changes elicited upon tryptophan omission from the diet are not mediated by *B. rodentium*. Altogether, these results point to the importance of multiple bacterial strains in tumor inhibition, and highlight the importance of tryptophan degradation in tumor growth control.

### B. uniformis phenocopies effects of B. rodentium

*In silico* genomic reconstruction of metabolic pathways relevant to the 11-species of microbes enriched in *Rnf5* KO mice revealed *B. rodentium* to be the only strain whose genome harbored the tryptophanase (*TnaA*) gene. As noted above, *TnaA* encodes an enzyme that degrades tryptophan. Given that others have linked tryptophan to inhibition of anti-tumor immunity^19,28,29^, we searched for *TnaA*-expressing bacterial species, particularly those found in the human gut. For these analyses we sought species or strains that also harbored *ArAT*, which encodes an enzyme also functioning in tryptophan degradation. In this effort, we identified *B. uniformis*, a species that expresses a *TnaA* orthologue and *ArAT* and is commonly found in the human gastrointestinal tract. When compared with other *Bacteroides* strains, *B. uniformis* is phenotypically and genetically close to *B. rodentium* (Supplemental Tables 4, 5). Administration of *B. uniformis* recapitulated changes observed with *B. rodentium* in inhibiting melanoma growth in GF mice as well as in mice harboring more complex gut microbiome, including GF mice colonized with a microbiota deficient in *Bacteroides* (MC608-F-a1), and in conventional C57BL/6J mice (Figures 4A, 4B, Supplemental Figure 4A, 4B). Notably, *B. uniformis* administration also inhibited growth of colorectal cancer, pancreatic cancer in MC608-F-a1 mice, and breast cancer in conventional Balb/c mice (Figure 4C). Notably, TCGA analysis revealed that the presence of *B. uniformis* was also associated with better remission in solid tumors (Figure 4D).

**Figure 4:**
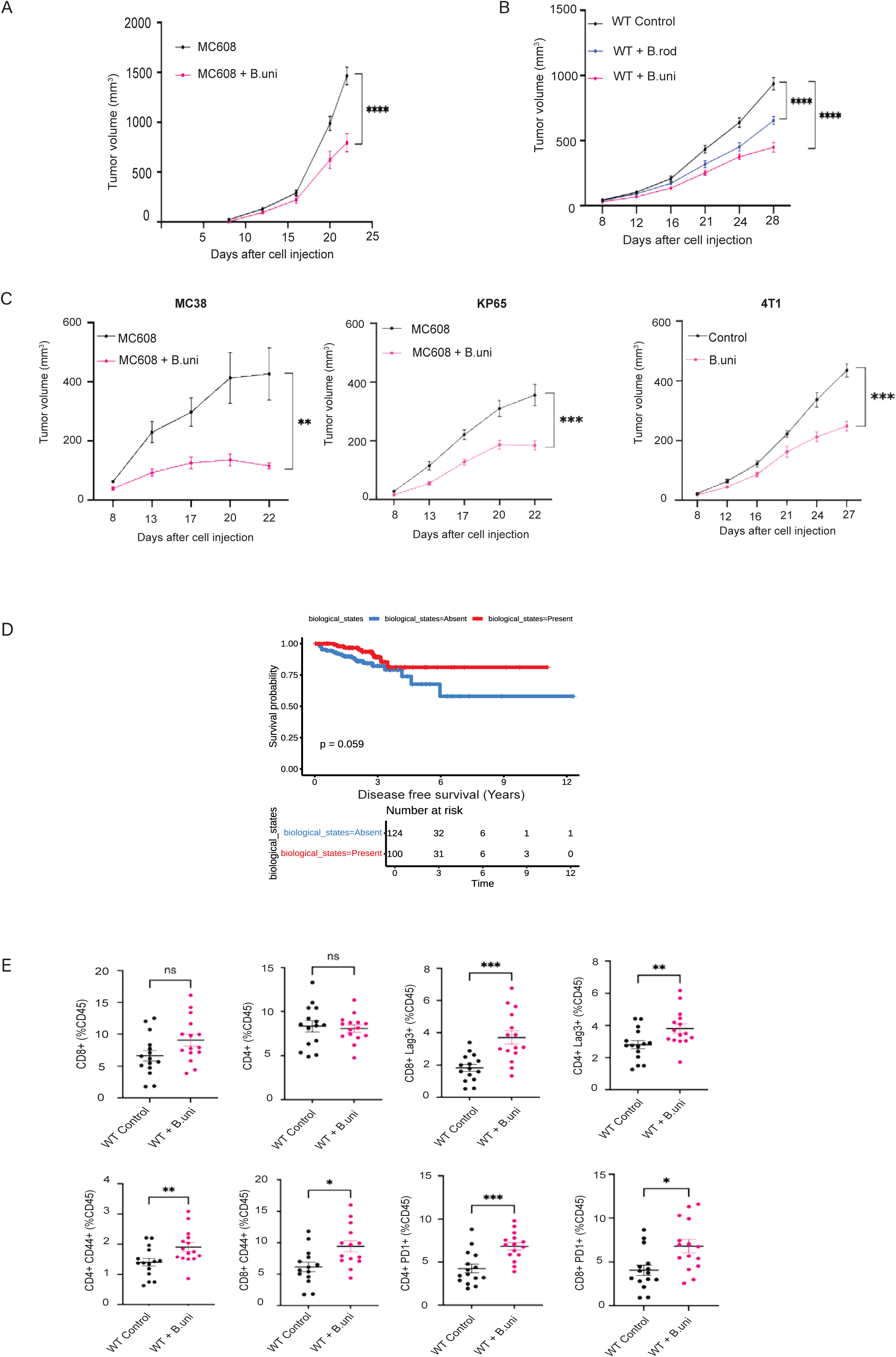
*B. uniformis* colonization phenocopied the tumor inhibition seen upon *B. rodentium* colonization. **(A)** Tumor growth in GF mice colonized with either microbiome MC608-F-a1 or MC608-F-a1 plus *B. uniformis via* oral gavage three times a week, starting 14 days prior to subcutaneous injection with YUMM1.5 cells (n = 8 mice/treatment). **(B)** Tumor growth in conventional C57BL/6J mice that were colonized with either *B. rodentium or B. uniformis via* oral gavage three times a week, starting 14 days prior to subcutaneous injection with YUMM1.5 cells (n = 7 mice/treatment). **(C)** Tumor growth in mice colonized with either MC608-F-a1 or MC608-F-a1 plus *B. uniformis via* oral gavage three times a week, starting 14 days prior to subcutaneous injection with MC38 cells or KP65 cells (n = 8 mice/treatment) and tumor growth in concentional Balb/c mice that were colonized with *B. uniformis via* oral gavage three times a week, starting 14 days prior to subcutaneous injection with 4T1 cells (n = 8 mice/treatment). **(D)** Kaplan-Meier analysis of % survival (disease-free survival, years) relative to the abundance of *B. uniformis.* Indicated are the biological states (presence or absence of the indicated bacterial species). The log-rank test p-value is indicated in the survival plot. Cox proportional hazards model results: HR = 0.50 (95% CI: 0.22–1.12), p = 0.09. **(E)** Quantification of tumor infiltration of CD8^+^ and CD4^+^ T cells, CD44^+^, Lag 3^+^ and PD1^+^ on CD4^+^ CD8^+^ T cell 12 days after injection of YUMM1.5 cells into concentional C57BL/6J mice that were colonized with *B. uniformis* (*via* oral gavage three times a week), starting 14 days prior to tumor inoculation until end of the experiment (n =15 mice/treatment). Data shown represent two experiments. Data were analyzed by unpaired t-test. *P < 0.05, **P < 0.005, ***P < 0.001, ****P < 0.0001 by two-tailed t-test or two-way ANOVA.

We next determined whether *B. uniformis* induces changes in factors or pathways associated with immune activity, phenotypes we have previously observed with *B. rodentium*. RT-qPCR analyses of RNA derived from melanoma tumors subjected to treatment with the indicated bacterial strains indicated that *B. uniformis* colonization promoted notable increases in expression of markers associated with pathogen-induced cytokine signaling pathways in CCR1-, TLR1- and TLR2-positive immune cells (Supplemental Figure 4C and 4D). We then used immunophenotyping to assess anti-tumor responses 12 days after melanoma tumor cell inoculation into conventional C57BL/6J mice. Fluorescence-activated cell sorting (FACS) of tumor-infiltrated immune cells on CD45-enriched cell populations and identified increased infiltration of CD44^+^ CD8^+^ and CD4^+^ T cells into tumors isolated from mice colonized with *B. uniformis* compared with controls in conventional C57BL/6J mice. These findings coincided with increased expression of the checkpoint receptors PD1 and Lag3 on CD8^+^ and CD4^+^ T cells (Figure 4E). We observed no changes in expression of markers associated with dendritic cells, macrophages or natural killer cells after *B. uniformis* administration (data not shown). Altogether, these results suggest that *B. uniformis,* like *B. rodentium*, promotes anti-tumor immunity.

### The enzyme encoded by TnaA gene is required for B. uniformis to inhibit melanoma growth

To confirm that tryptophan degradation is required for anti-tumor immunity promoted by *B. uniformis,* we deleted *TnaA* gene by allelic exchange, that was performed in the *B. uniformis* genome. PCR was carried out to confirm the deletion of *TnaA,* compared WT form of *B. uniformis* (Supplemental Figure 5A). In agreement, indole production, which was monitored *in vitro* in cultures of the bacterial secretome of either *TnaA* mutant or WT forms of *B. uniformis* revealed a significant decrease by the *TnaA* mutant, compared to the WT *B. uniformis* strain (Supplemental Figure 5B). Notably, while colonization with WT *B. uniformis* inhibited melanoma tumor growth in GF and ASF-bearing GF mice, this protective effect was absent in mice colonized with *TnaA* deleted *B. uniformis*. These findings overall strongly suggest that tryptophan degradation by a single gut bacterial strain is required to inhibit melanoma growth (Figure 5A, 5B, Supplemental Figure 5C, 5D). To further confirm the loss of indole production we perform metabolomic analysis of indoles in cecal samples from mice that were mono-colonized with either the WT or TnaA deleted forms of *B. uniformis*. This analysis confirmed that mice colonized with WT but not *TnaA* mutant form of *B. uniformis* produced indole (Supplemental Figure 5E).

**Figure 5:**
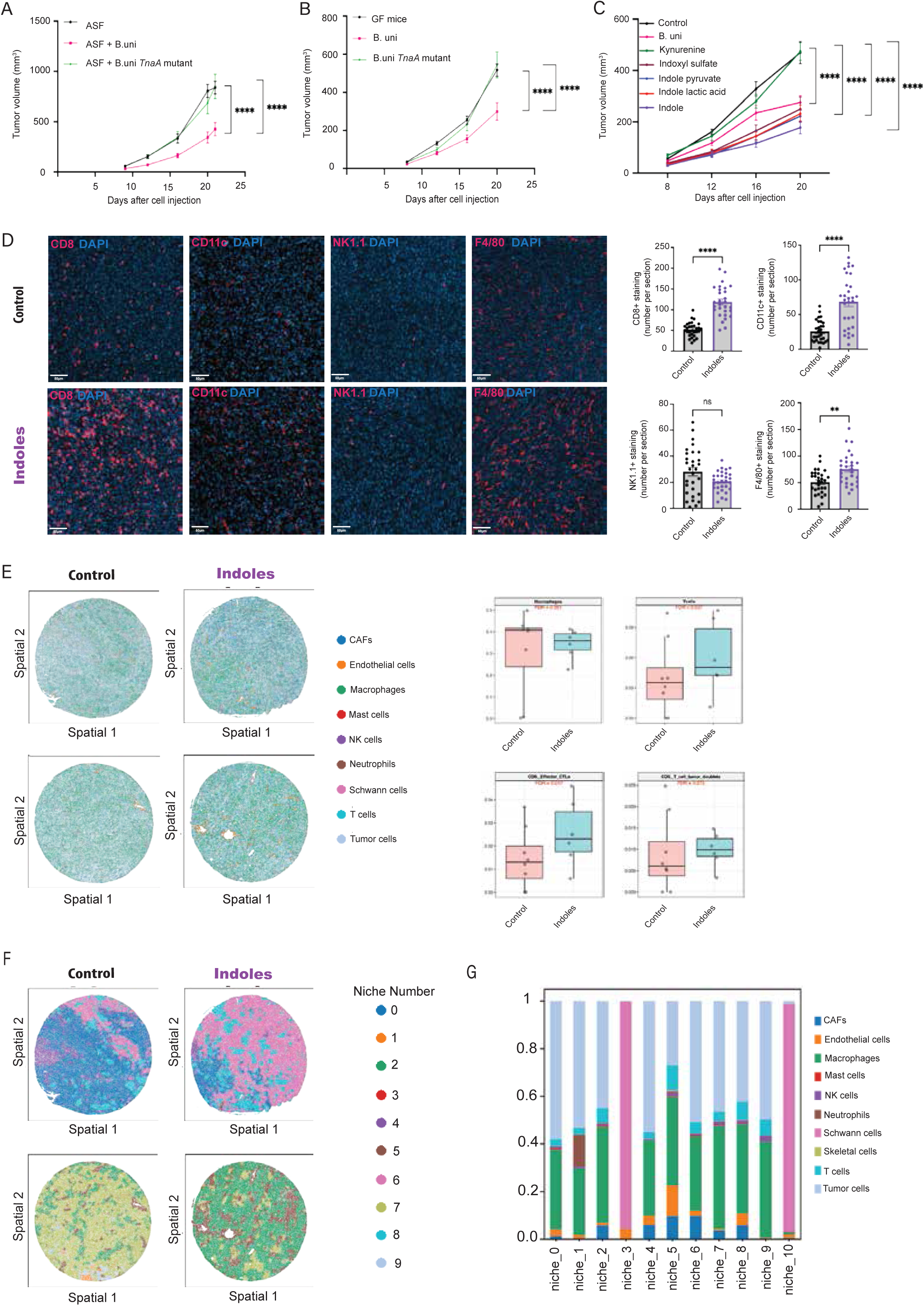
*B. uniformis* colonization phenocopied tumor inhibition seen upon *B. rodentium* colonization requires the tryptophanase (*TnaA*) gene. **(A)** Tumor growth in GF mice colonized with either ASF, ASF plus *B. uniformis,* or ASF plus *B. uniformis* tryptophanase mutant 14 days prior to YUMM1.5 tumor cell injection (n = 15 mice/treatment). **(B)** Tumor growth in GF mice colonized with either *B. uniformis,* or *B. uniformis TnaA* mutant 14 days prior to YUMM1.5 tumor cell injection (n = 15 mice/treatment). **(C)** Tumor growth in conventional C57BL/6J mice that were treated via oral gavage daily, starting one day after subcutaneous injection of YUMM1.5 cells. Mice were treated with kynurenine, indoxyl sulfate, indole pyruvate, indole lactic acid, and indole (n=6 mice/treatment). **(D)** Staining with anti-mouse CD8, CD11c, NK1.1, and F4/80 antibodies of tumor sections (Control, n=3; Indoles, n=3). CD8^+^, CD11c^+^, NK1.1^+^,and F4/80^+^ positive cells were quantified. For image analysis, 10 images were randomly selected from different areas of each tumor, in which the number of positive cells per section were counted. **(E)** Spatial distribution of immune cell types in tumor samples and their quantification (Control, n=3; Indoles, n=3). **(F)** Representative images of spatial distribution for niches enrichment. (Control, n=3; Indoles, n=3). **(G)** Bar chart showing the composition of the cell types in different niches. Data shown represent two experiments. Data were analyzed by unpaired t-test. *P < 0.05, **P < 0.005, ***P < 0.001, ****P < 0.0001 by two-tailed t-test or two-way ANOVA.

We next analyzed cytokine levels in sera of GF mice colonized with ASF plus *B. rodentium* compared with ASF only controls. That analysis revealed that GF administered with *B. rodentium* plus ASF showed lower levels of G-CSF and CXCL1 than did GF mice harboring only the ASF (Supplemental Figure 5F). Similarly, G-CSF and CXCL1 levels were lower in sera from both ASF-bearing GF and conventional C57BL/6J mice administered either *B. rodentium* or *B. uniformis* than in sera from mice colonized with *TnaA* mutant *B. uniformis* (Supplemental Figure 5F). Both G-CSF and CXCL1 (also known as Keratinocyte Chemoattractant; KC) cytokines have been previously implicated in inhibition of anti-tumor immunity^30–33^. These findings suggest that G-CSF and CXCL1 inhibition by either *B. rodentium* or *B. uniformis* may serve to promote the activation of factors associated with immune activation against melanoma.

### Indoles are sufficient to inhibit melanoma growth in mice

We next asked whether tryptophan degradation products such as indoles could phenocopy anti-tumor effects of *B. uniformis* or *B. rodentium*. To do so, we administered metabolites produced by either *TnaA* or *ArAT* enzymes, which are encoded by these bacterial strains, by daily oral gavage to conventional C57BL/6J mice. The degree of tumor inhibition was then compared to that of mice administered metabolites, which are produced by host enzymes upon tryptophan degradation and are linked to IDO pathway. Notably, we found that metabolites such as indole and indoxyl sulfate (both *TnaA* breakdown products) as well as indole lactic acid and indole pyruvate (*AraT* breakdown products) effectively inhibited melanoma growth in these mice. Conversely, kynurenine, a metabolite produced by the host *via* the IDO pathway, did not inhibit tumor growth (Figure 5C and Supplemental 5G).

We next determined whether administration of indoles induces anti-tumor immunity as was observed following the administration of *B. rodentium* and *B. uniformis*. To this end we performed RT-qPCR analyses of select genes in melanoma tumors from mice subjected to treatment with indoles, compared with controls. Notably, indoles promoted an increase in the expression of CD8, CD4, CD3, and NK1 (Supplemental Figure 5H). Further analysis revealed an increase in the infiltration of CD8^+^ T cells, CD11c^+^ cells, and F4/80^+^ cells into tumors obtained from mice treated with indoles compared to controls (Figure 5D). Immunophenotyping using FACS, performed 12 days after melanoma tumor cell inoculation into conventional C57BL/6J mice, identified increased infiltration of CD44^+^, CD69^+^, and CD62L^+^ on CD8^+^ T cells from CD45-enriched cell populations. Increase expression of GZMB, TNFα, and IFNγ was noted on CD8^+^ and CD4^+^ T cells isolated from tumors of mice treated with indoles compared with controls, in conventional C57BL/6J mice. These findings coincided with increased expression of the checkpoint receptor PD1 on CD8^+^ T cells (Supplemental Figure 5I). Independent analyses were performed to determine the spatial distribution of the immune cells within tumor. Confirming earlier observations we identified that tumor that were obtained from mice that were treated with indoles exhibited increased infiltration of T cells populations, among which the infiltration of CD8^+^ effector cytotoxic T cells, and macrophages populations were most pronounced (Figure 5E). Evaluation of different niche populations revealed that tumors that were obtained from indoles treated mice exhibited increase in niches associated with the presence of CAFs, macrophages and T cells (niches 2,6; Figures 5F and 5G). Collectively, these findings confirm that indoles promote anti-tumor immunity.

As indoles serve as ligands to activate the transcription factor AhR^34,35^, we asked whether AhR signaling is involved in melanoma inhibition induced by indoles. To this end, we monitored changes in the expression of AhR signaling components by RT-qPCR in tumors from mice subjected to indole treatment compared with controls. This analysis identified increased expression of CyP1B1 genes and decreases in expression of TIPARP, BAFF and CyP1A1 (Supplemental Figures 6A and 6B). To further assess whether AhR signaling pathways may be involved in melanoma growth inhibition induced upon *B. rodentium* inoculation, we analyze the effect of the AhR agonist TCCB *in vivo*. While GF mice colonized with ASF plus *B. rodentium* inhibited melanoma growth, the addition of TCCB to this setting abolished this inhibition (Supplemental Figure 6C). These data implies that AhR may not mediate tumor growth inhibition seen upon *B. rodentium*, or indole administration. As indoles also can activate PXR and GPCR signaling pathways, we perform RT-qPCR of components associated with these pathways. Indoles administration did not increase expression of components associated with PXR (ABCB1, ZO-1 and NF-κB) or GPCR (FOS, EGR1, JUN, NR4a1, IL-6; Supplemental Figures 6D, 6E). These observations suggest that signaling pathways other than AhR, GPCR and PXR may mediate indole ability to induce anti-tumor immunity, an important aspect that will be elucidated in future studies.

### ArAT and TnaA expression increases in patients who respond to ICB

Given that immune checkpoint therapy is currently a first line therapy for melanoma patients, we analyse stool samples from different human cohorts of melanoma patients, comparing responders versus non-responders to ICB therapy, to identify possible differences in the level of enzymes implicated in tryptophan degradation. This analysis identified a significant increase in the level of *ArAT* in patients who respond to ICB therapies in one of the cohorts (Figure 6A) and an increase in *TnaA* levels in Gunjur^71^ and Lee^70^ Cohorts (Figure 6B). In addition, we analyze dysbiosis levels in these cohorts, and the *TnaA* levels were found to be elevated in patients who are eubiotic compared with those who have dysbiosis (Figure 6C). In agreement, increase in metabolites or enzymes implicated in tryptophan degradation pathways was independently observed in patients who responded to ICB therapies in different cancer types^36^, such as melanoma^37^, NSCLC^38,39^, and CRC^40^. These findings establish the human relevance of our finding that tryptophan degradation is implicated in the inhibition of melanoma and identifies indoles, produced by either *B. rodentium* or *B. uniformis,* as important metabolites sufficient to mediate inhibition of melanoma growth.

**Figure 6:**
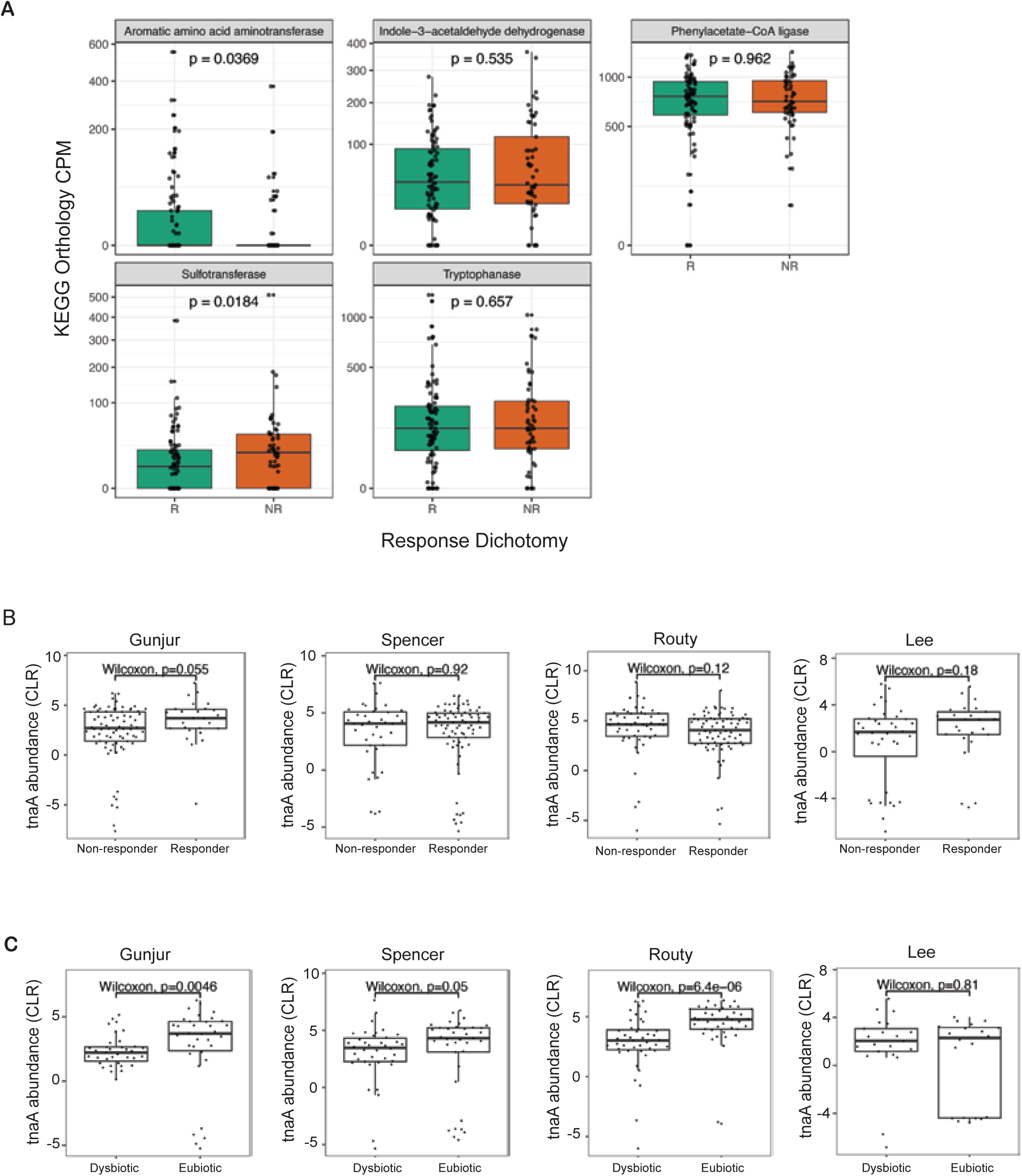
*ArAT* and *TnaA expression increases in patients who respond to ICB.* **(A)** Each panel displays gene-level abundance (counts per million, CPM) for the indicated KEGG orthology, stratified by response dichotomy (R, responders; NR, non-responders). Boxes represent the interquartile range with the median line; whiskers extend to 1.5× IQR. Statistical comparisons performed using the Wilcoxon rank-sum test; p-values are shown above each comparison. **(B)** Abundance of *TnaA* in Ganjur^71^, Spencer^72^, Routy^49^, and Lee^70^ cohorts, comparing responder *vs.* non-responders to immune checkpoint therapy **(C)** Abundance of *TnaA* in Ganjur^71^, Spencer^72^, Routy^49^, and Lee^70^ cohorts. Functional dysbiosis was computed for all the samples (described in^69^). Samples were classified, per study, as dysbiotic (top tertile of functional dysbiosis) or eubiotic (bottom tertile of functional dysbiosis) levels.

## Discussion

Here, we identify select bacterial species and their respective metabolites as capable of limiting melanoma growth. Both the mouse and human strains (*B. rodentium* and *B. uniformis*, respectively) were shown capable of inhibiting melanoma, as well as other tumors, in different cancer models and mouse strains. This inhibition required the bacterial enzymes *TnaA* and *ArAT* which are important in tryptophan breakdown to indoles. While mutation of *TnaA* in *B. uniformis* was sufficient to abolish tumor inhibition phenotype, elevated level of *ArAT* was found in patients who responded to ICB.

While our initial findings were based on colonization of select bacteria in GF mice, subsequent experiments performed using mouse models with more complex microbiomes, including GF mice colonized with a microbiota devoid of *Bacteroides* species and conventional C57BL/6J mice, all of which showed anti-tumor effects resembling those seen in GF mice. We found that changes elicited by tryptophan breakdown were linked to activation of intestinal immune cells, inhibition of serum factors thought to limit antitumor immunity (CXCL1 and G-CSF) and tumor infiltrations by CD8^+^ T cells, activities coinciding with enhanced anti-tumor immunity.

The importance of the gut microbiota in controlling anti-tumor immunity and tumor growth has been intensively investigated in several tumor models^3,41–44^. In mouse models of pancreatic cancer, indoles produced in the gut by *Lactobacillus* species reduced anti-tumor immunity by activating the aryl hydrocarbon receptor (AhR) in tumor cells and thereby decreasing CD8^+^ T cell levels and increasing tumor growth^19^. In contrast, the metabolite indole-3-aldehyde (I3A) produced by *Limosilactobacillus reuteri* increased chemotherapy efficacy in mouse models of pancreatic cancer, and patients with pancreatic cancer who responded to therapy showed enrichment of *Lactobacillus reuteri*^45^ in the gut. I3A has also been linked to enhanced immune checkpoint inhibitor efficacy against melanoma in mouse models and in patients^37^. Conversely, indole-3-lactic acid, which is produced by *Lactobacillus gallinarum*, has been shown to limit colorectal cancer by inhibiting AhR expression in tumor cells^46^. While a number of AhR ligands were shown to impact degree of anti-tumor immunity, future studies will clarify which of the cellular signaling components mediate *B. rodentium* / *B. uniformis* or indoles ability to induce anti-tumor immunity. Importantly, we observed that *B. uniformis* colonization also inhibited melanoma growth, phenocopying effects of *B. rodentium*. Both of these species harbor genes encoding the tryptophan-degrading enzymes *TnaA* and *ArAT*, which produce different types of indoles as degradation products. Notably, each of the indoles tested here for anti-tumor activity, whether produced by *TnaA* or *ArAT*, was sufficient to inhibit melanoma growth. These findings coincide with human data showing that elevated *ArAT* expression is seen in melanoma patients that respond to immune checkpoint blockade. Along these lines, the presence of *B. uniformis* has been linked with remission of cancer patients^47,48^. Along these lines, metataxonomic microbiome analyses of responders to PD1 therapy, also pointed to a positive correlation between treatment responsiveness and *B. uniformis* abundance^49^. Overall, our findings suggest that indoles produced by enzymes presence in *B. uniformis* are critical for tumor growth inhibition, an effect shown here for melanoma, breast, colon and pancreatic cancers, indicative of a general phenomenon across different genetic mouse models.

Independent studies suggest that the mechanisms underpinning *B. uniformis* anti-tumor activity may involve the gut-brain axis. In rats, colonization of *B. uniformis* CECT 7771 has been reported to impact brain circuitry functioning in the reward response^50^. Given that the brain reward system is implicated in enhancing anti-tumor immunity and limiting tumor growth in mice^51^, activity of the gut-brain axis^52^ may also underlie effects of *B. uniformis* in our model. *B. uniformis* has also been implicated in attenuating intestinal inflammation as its administration restored intestinal barrier function, increased NF-κB activity and mitogen-activated protein kinase (MAPK) signaling in colonic tissues, and promoted TH17 cell differentiation in a mouse model of checmically induced colitis^53^. Such findings suggest that enhancing anti-tumor immunity *via B. uniformis* may also impact auto-immune responses, which often emerge in response to immune checkpoint therapies^54,55^, exemplified in the *Rnf5* KO mice, which exhibit both enhanced anti-tumor immunity and autoimmunity^4,11^.

In all, this study highlights a novel approach for eliciting effective anti-tumor immunity to inhibit tumor growth, an activity that requires indole production due to degradation of tryptophan by *Bacteroides* species in the mouse and human intestine. How indoles impact anti-tumor immunity remains to be determined, our data suggest the canonical transcriptional pathways associated with AhR, GPCR or PXR activation are not strongly induced following *B. rodentium* or indoles administration, pointing to the requirement to identify the signaling components that mediate anti-tumor immunity phenotypes, a subject for independent future studies. Notably, a partial, context-dependent or cell type specific AhR/GPCR/PXR signaling cannot be excluded. The finding that indoles serve as key mediators of anti-tumor phenotypes may offer alternative paths for clinical evaluation of the tryptophan degradation pathway, distinct from previously attempted approaches to inhibit IDO1. Our findings provide the foundation for treatment modalities that could complement or replace existing ones.

## Supporting information

RONAI SUPP FIG

## STAR★Methods

### Key resources table and detailed methods

### Animals and tumor model

All experimental animal procedures were approved by the Institutional Animal Care and Use Committee of Sanford Burnham Prebys Medical Discovery Institute (SBP: approval #22-080) and Cedars-Sinai Medical Center (Protocol: IPROTO202400000337) and complied with respective ethical animal testing and research regulations. Male mice, 6-8 weeks-old, were used for all experiments. Conventional C57BL/6J mice, C3H, and Balb/c mice were purchased from Jackson Laboratories. GF C57BL/6J mice were bred and maintained at the University of Nebraska-Lincoln (UNL) by the Nebraska Gnotobiotic Mouse Program under gnotobiotic conditions in flexible film isolators. The Institutional Animal Care and Use Committee at UNL approved experiments involving GF mice (protocols #2126 and #2622). All GF mice were fed autoclaved chow diet ad libitum (LabDiet 5K67, Purina Foods). GF status of the breeding colony was routinely checked as described^56^. Mice were injected subcutaneously (s.c.) with either 1×10^6^ YUMM1.5 melanoma cells, 8×10^5^ SW1, melanoma cells, 5×10^5^ 4T1 breast tumor cells, 2×10^5^ KP65, pancreatic cells or 5×10^5^ MC38, colorectal cancer cells. Tumor size was measured twice a week to calculate tumor volume. Tumors were weighed at the time of excision.

### Cell lines

This study used the melanoma line YUMM1.5^57^ and SW1, colorectal line MC38, pancreatic cell line KP65 and breast cancer cell line 4T1, cultured in Dulbecco’s modified Eagle’s medium supplemented with 10% fetal bovine serum (FBS) and antibiotics. Cells were authenticated and determined to be mycoplasma-free.

### Culture and administration of bacterial strains

Two microbiomes—the Altered Schaedler Flora (ASF) and MC608-F-a1—were used to inoculate GF mice. The ASF consists of eight strains: ASF 356, *Clostridium* sp.; ASF 360, *Lactobacillus intestinalis*; ASF 361, *Lactobacillus murinus*; ASF 457, *Mucispirillum schaedleri*; ASF 492, *Eubacterium plexicaudatum*; ASF 500, *Pseudoflavonifractor* sp.; ASF 502, *Clostridium* sp.; and ASF 519, *Parabacteroides goldsteinii*. MC608-F-a1 refers to mice derived from a wild mouse population in the Massif Central region of France previously maintained at the Max Planck Institute for Evolutionary Biology (Plon, Germany; referred to as MC608-F-a1 in ^22,23^ and as A in ^24^). Ceca were collected from either ASF-or MC608-F-a1-bearing mice, stored at −80°C, and the contents resuspended (1:3 wt/vol) in reduced PBS as previously described ^58^ at the time of use.

Two weeks before tumor cell injection, 100 µL of cecal contents (either from ASF- or MC608-Fa1-bearing mice) and bacterial strains of interest (1×10^6^ to 1×10^7^) were administered to GF mice via three oral gavages one day apart. Bacterial strains used in this study included: *Bacteroides rodentium* ST28 (DSMZ 26882), *Bacteroides uniformis* VPI 0061 (ATCC 8492), *Phocaeicola massiliensis* B84634 (DSM 17679), and *Phocaeicola sartorii* AC20 (DSM 21941). For experiments performed in conventional C57BL/6J mice, *B. rodentium* and *B. uniformis* were colonized via oral gavage three times a week for the duration of the study.

### Deletion of the *B. uniformis tnaA* gene by allelic exchange

A pLGB13-based^59^ gene deletion plasmid (pLGB13-ΔtnaA) for deleting tnaA was constructed using Phusion High-Fidelity DNA Polymerase for all PCR steps. Briefly, the upstream and downstream regions (∼1500bp) of the *tnaA* gene were amplified from the *Bacteroides uniformis* genome using the following two primer sets: (upstream: TAAATCTCTTACATAGACTCTGTATCTTTAATAATTTGACA and ACTGGAAGATAGGCAATTAGGAAGAAGAGCAAGTACGGCTG) and (downstream: GTAAGATTAGCATTATGAGTGGTTCCACCTCTTTCTTGAACAAGC and CAGAGTCTATGTAAGAGATTTACAGACTCAATGTCA), respectively. PCR products were visualized on a 1% agarose gel, and fragments of the expected size were gel purified using the QIAquick Gel Extraction Kit (Qiagen). The deletion vector pLGB13 was linearized by double digestion with BamHI and SalI restriction enzymes (New England Biolabs). Digested pLGB13 and the gel-purified upstream and downstream *tnaA* fragments were then ligated using a Gibson Assembly Master Mix (New England Biolabs), following the manufacturer’s protocol. The assembled plasmid was transformed into electrocompetent *E. coli* strain S17-λ*pir* and transformants were selected on LB agar plates supplemented with 200 μg/mL ampicillin. Successful insertion of the upstream and downstream regions was confirmed by PCR using primers that lie inside the 1.5 kb fragment used to create pLGB13-ΔtnaA (GTTTTCGTTGTTCACGATGGTT/ and CTTGTCGGTCAGCTCCAAC). The entire pLGB13-ΔtnaA construct was then sequenced by Plasmidsaurus to verify the absence of off-target mutations and confirm the correct arrangement of upstream and downstream region.

The pLGB13-ΔtnaA plasmid in conjugative *E. coli* donor strain S17-λpir was used to construct the gene deletion. For conjugation, both the donor *E. coli* S17-λpir and the recipient *B. uniformis* were grown overnight. *E. coli* was cultured in LB broth supplemented with 200 μg/mL ampicillin at 37°C with shaking, while *B. uniformis* was cultured in TYG broth at 37°C under anaerobic conditions. On the day of conjugation, overnight cultures were subcultured (1:500 for *E.coli* and 1:100 for *B. uniformis*) into fresh respective media. Cells were incubated until they reached OD_600_ of approximately 0.1-0.2. Donor and recipient cultures were pelleted by centrifugation at 20,000 x g for 3 minutes and washed twice with sterile TYG to remove residual antibiotics. The donor and recipient cells were then resuspended in 200 μl of fresh TYG. Four 50 μl spots of the mixed bacterial suspension were then spotted directly onto pre-warmed Blood BHI agar plates and incubated aerobically at 37°C overnight. Following incubation, bacteria from the mating spots were resuspended in 1 mL of sterile TYG. Serial dilutions of this suspension were then plated onto selective Blood BHI agar plates containing 25 μg/ml erythromycin and 200 μg/ml gentamicin. Plates were incubated anaerobically at 37°C for 2 to 3 days. Individual colonies (n=12) were re-streaked for isolation on fresh selective plates to ensure purity.

Twelve isolated merodiploids (erythromycin resistant colonies containing the *tnaA* deletion plasmid integrated via one of the cloned gene-flanking fragments) were individually grown overnight in 5 mL of TYG broth. These cultures were then mixed in equal volumes, serially diluted, and plated onto BHI agar blood agar plates containing 100 ng/mL anhydrotetracycline (aTC). Plates were incubated anaerobically at 37°C for 2 to 4 days. Single colonies that grew on aTC-containing plates were re-streaked for isolation. Loss of the target *tnaA* gene was confirmed by genomic PCR using the upstream and downstream nested primers GTTTTCGTTGTTCACGATGGTT and CTTGTCGGTCAGCTCCAAC, respectively, and testing in medium containing erythromycin to confirm that the plasmid was also lost from the strain (*i.e*., clones had regained sensitivity to this antibiotic).

### Dietary manipulations

For dietary experiments, we used irradiated chow, either Amino Acid diet (TD.01084, Envigo) or Tryptophan Deficient AA Diet (TD.180506, Envigo). Diets were administered to mice starting 3 days before tumor inoculation until the end of the experiment.

### Taurine, kynurenic acid, kynurenine, and indoles administration

Taurine (10 mg/mL, Sigma Aldrich) was administered in drinking water 3 weeks before tumor cell injection until the end of the experiment. Sterile water served as vehicle control. Kynurenic acid (Sigma Aldrich; 40 mg/kg body weight) in 200 µL sterile water was administered daily by oral gavage starting 3 days before tumor cell inoculation until the end of the experiment. Kynurenine, indoxyl sulfate, indole pyruvate, indole 3 lactic acid and indole (All from Sigma Aldrich; 40mg/kg body weight) in 100 μL of corn oil was administered daily by oral gavage starting 1 day after tumor cell inoculation until the end of the experiment.

### In vivo antibody treatment

For anti-PD1 therapy mice were administered anti-PD1 (clone RMP1-14) or the control (rat IgG2a isotype) antibodies on days 7, 10, 13, 16, 18, 22, 25 after tumor cell inoculation (200μg administered by I.P. injection). For inhibiting IDO1, Epacadostat (MedChemExpress) was administered (oral gavage, 60μg/kg) daily, starting at day 7 after tumor inoculation until the end of the experiment. TCDD (Cambridge Isotope Laboratories), was administrated (oral gavage 5ug/kg) weekly, starting two weeks prior tumor injection.

### RNA extraction and RT-qPCR analyses

Total RNA was extracted from mouse tumor samples using the Total RNA Miniprep Kit (Monarch). RNA purity and concentration was quantified by reading at 260 and 280 nm in a NanoDrop spectrophotometer (Thermo Fisher). RNA was reverse transcribed using a High-Capacity Reverse Transcriptase Kit (Invitrogen) according to the manufacturer’s protocol. RT-qPCR analysis was performed using SsoAdvanced Universal SYBR Green Supermix (Bio-Rad) and a Bio-Rad CFX Connect Real-Time system. Expression levels were normalized to H3N controls. The sequences of the primers used are listed in Supplemental Table 1.

### Gas Chromatography-Mass Spectrometry (GC-MS) Sample Preparation and Analysis

Samples of cecum contents (40-60 mg) or serum (15 µl) and standards were prepared as described ^60^, with additional standards as listed in Supplemental Table 2. Samples and standards were analyzed by GC-MS using an TG-SQC column (15 m x 0.25 mm x 0.25 µm, Thermo) installed in a Thermo Scientific TSQ 9610 GC-MS/MS. The GC was programmed with an injection temperature of 300°C and a 0.5 µl injection. The GC oven temperature was initially 130°C for 4 min, rising to 250°C at a rate of 6°C/min, and to 310°C (280°C for serum) at 60°C/min with a hold at the final temperature for 2 min. GC flow rate with helium carrier gas was 50 cm/s. The GC-MS interface temperature was 300°C and (electron impact) ion source temperature was 200°C, with 70 eV ionization voltage. Standards were run in parallel with samples. Metabolites in samples and standards were detected by MS/MS using precursor and product ion masses, and collision energies shown in the attached table. Sample metabolites were quantified using calibration curves made from the standards in Themo Chromeleon software, and further data processing to adjust for the relative quantities of metabolites in the standards, and for recovery of the internal standard were done in MS Excel.

Independent metabolomic analysis was performed at Metware Biotechnology Inc. Samples stored at −80°C were thawed on ice. 400uL solution (Methanol: water = 7.3, V/V) containing internal standard were mixed with 20mg of the samples (vortexed for 3 min). Samples were than sonicated (ice bath for 10 min) and vortexed (1 min) followed by incubation (−20°C) for 30min. The samples were centrifugated (12000 rpm for 10 min) and the supernatant was subject for another centrifugation (12000rpm for 3min). 200uL of the supernatant was than used for LC-MS analysis. For untargeted detection, Ultra Performance Liquid Chromatography (UPLC) and Quadrupole-Time of Flight Spectrometry with a chromatographic column, ACQUITY HSS T3 (2.1 x 100mm, 1.8 µm) were used using ultrapure water (0.1% formic acid added), and acetonitrile (0.1% formic acid added) with column (temperature 40°C, flow rate 0.4ml/min, and injection volume of 5 μl). For widely targeted detection, UPLC and tandem mass spectrometry (MS/MS) (QTRAPr6500+) were employed using chromatographic column (Waters ACQUITY UPLC HSS T3 C18 1.8um, 2.1mm*100mm). A phase was ultrapure water (0.1% formic acid added), B phase was acetonitrile (0.1% formic acid added); Gradient program was 95:5 V/V at 0 min, 10:90 V/V at 10,0min, 10:90 V/V at 11.0min, 95:5 V/V at 11.1min, 95:5 V/V at 14.0min; Flow rate was 0.4mL/min at 40 °C, injection volume 2uL.

### RNAseq analysis

RNA samples were prepared from mouse tumor samples from ASF alone and ASF plus *B. rodentium* samples collected at experiment’s end (Day 22). Samples contained 250 ng of RNA. For library construction, PolyA RNA isolation and library preparation were performed with the Watchmaker Genomics mRNA Library Prep Kit (Watchmaker Genomics) and Elevate Long UDI Adapters (Element Biosciences). RNA libraries were pooled and sequenced (2×75bp) on the Element Biosciences AVITI sequencer with the 2×75 Cloudbreak kit.

Illumina Truseq adapters and polyA/polyT sequences were trimmed from raw reads using Cutadapt v2.3^61^, and trimmed reads were aligned to mouse genome version mm10 and Ensembl gene annotations v84 using STAR version 2.7.0d_0221^62^, adopting alignment parameters from the ENCODE long RNA-seq pipeline (https://github.com/ENCODE-DCC/long-rna-seq-pipeline). RSEMv1.3.1 was used to obtain gene level estimated counts and transcripts per million^63^.

FastQC v0.11.5 (https://www.bioinformatics.babraham.ac.uk/projects/fastqc/) and MultiQC v1.8^64^ were used to assess quality of trimmed raw reads and alignment to the genome and transcriptome. Only genes with RSEM estimated counts ≥5 times the total number of samples were retained for differential expression analysis. Differential expression comparisons were performed using the Waldtest implemented in DESeq2 v1.22.2.^65^ Genes with a Benjamini– Hochberg-corrected P-value of< 0.05 and fold-change of ≥1.5 or ≤1.5 were defined as differentially expressed. Gene set enrichment analysis (GSEA) for TCGA-PRAD comparison of ASF versus ASF plus *B. rodentium* was performed using the pre-ranked option in GSEA version 4.2.3(RRID:SCR_003199)^66^. GSEA for RNA-seq comparisons was performed using GSEA version 4.3.2, and RPKM values with the parameter “Permutation type = gene_set” Pathway analysis was performed using Ingenuity Pathway Analysis (IPA) software (Qiagen).

### Histology and immunofluorescence

Tumor and small intestine samples were collected immediately after animals were sacrificed. The small intestine was split open lengthwise, rinsed, and rolled up from the proximal to distal end to form a “Swiss roll.” Tissues were fixed in 4% formalin, washed with PBS, embedded in paraffin, and cut into 5 µM-thick sections and stained with H&E.

For immunohistochemistry, tissue sections were deparaffinized, rehydrated and washed with PBS. Antigen retrieval was performed using Epitope Retrieval Solution, pH 6.0 (Leica Biosystems). Sections were incubated overnight at 4°C with CD8a (Cell signaling technology, D4W2Z, dilution: 1:500), CD11c (Cell signaling technology, D1V9Y, dil: 1:200), F4/80 (Cell signaling technology, D4C8V, dilution: 1:500), NK1(Cell signaling technology, E6Y9G, dilution: 1:150), in Dako antibody diluent. Slides were then washed three times with PBS and incubated for 2h at room temperature with Alexa Fluor 594-conjugated and Alexa Fluor 488-conjugated secondary antibodies. Nuclei were stained with SlowFade Gold Antifade reagent (Vector) with 4’,6-diamidino-2-phenylindole (DAPI, Vector). Data were obtained using Olympus TH4-100 and Leica DMi8 Fluorescence microscopes.

### Tumor digestion

Tumors were harvested at day 12 after tumor injection and digested with 100 μg/mL DNase I (Sigma) and 1mL of collagenase D (Roche) at 37°C for 1 hour. To generate a single-cell suspension, the digested material was passed through a 70μm cell strainer. The cells were then washed two times with phosphate-buffered saline (PBS) and incubated with the indicated antibodies for flow cytometry.

### Flow Cytometry

Tumor single cells were washed with FACS staining buffer, before incubated with the indicated antibodies (dilution 1:200) at 4°C for 20 min. Cells were then fixed using 1% of formaldehyde in PBS (for 20 min), followed by two washes before resuspended in FACS staining buffer. The following antibodies were used: NK1.1 (Clone: OK136), CD25 (Clone: 3C7), CD8 (Clone: 53-6.7), CD4 (Clone: RM4-5), Lag3 (Clone: C9B7W), CD69 (Clone: H1.2F3), B220 (Clone: RA3-6B2), CD44 (Clone: IM7), CD45 (Clone: 104), CD279 (Clone: 29F.1A12), MHC II (Clone: M5/114.15.2), CD11b (Clone: M1/70), PDCA-1 (Clone: 129c1), CD11c (Clone: N418), MHC I (Clone: AF6-88.5), F4/80 (Clone: BM8), CD206 (Clone: C068C2), GR1 (Clone: RB6-8C5), CD80 (Clone: 16-10A1). All the data were collected on an BD FACSymphony A5 and analyzed using Flowjo Software (10.8.2).

### Spatial processing

#### Processing of samples

Formalin fixed paraffin embedded Tissue Micro Array (TMA) were used. Mouse sample were profiled using in situ Xenium platform (10x Genomics). Samples were mounted onto the Xenium slides, followed by fixation of the samples according to the manufacturer’s protocol (Xenium in Situ - FFPE Tissue Preparation CG000578 | Rev F). Target transcripts were hybridized with Xenium Mouse Tissue Atlassing predesigned Gene expression probes (Xenium in Situ Gene Expression CG000749 | Rev B). After probe hybridization, ligation and amplification were performed, and the tissues were stained with cell segmentation markers overnight, followed by nuclear staining (DAPI). The slides were then loaded onto the Xenium Analyzer. Following imaging, the slides were stained for H&E (Xenium Post-Xenium Analyzer H&E Staining CG000613 | B).

#### TMA core extraction and cell segmentation

Whole-slide DAPI images were loaded into QuPath (v0.6.0), where rectangular regions of interest (ROIs) were manually annotated around each TMA core. Core boundary coordinates (in microns) were exported using a custom Groovy script that converts pixel coordinates to physical units using the slide’s pixel calibration. Raw Xenium transcript tables were spatially subset per core using these coordinate boundaries. Transcripts with quality value (QV) < 20 were excluded, along with BLANK, NegControl, and UnassignedCodeword probes, yielding 183.7 million quality-filtered transcripts across the 14 cores. Cell segmentation was performed using Baysor(v0.7.1) on per-core transcript CSV files with Xenium-assigned cell IDs provided as prior segmentation labels.

#### Quality control and processing

Baysor outputs were loaded and concatenated into a single AnnData object. Quality filtering removed cells with fewer than 5 transcripts or fewer than 3 unique genes, retaining 504,862 cells across 379 genes from an initial 506,105 Baysor-segmented cells (1,243 cells removed). Raw transcript counts were stored in a separate layer, and expression values were normalized to counts per 10,000 followed by log1p transformation.

#### Batch integration and dimensionality reduction with scVI

Batch effects across slides were corrected using scVI (scvi-tools v1.3.0). A variational autoencoder was trained with the following parameters: 10 latent dimensions, 128 hidden units, 1 encoder/decoder layer, zero-inflated negative binomial (ZINB) gene likelihood, maximum KL divergence weight of 1.0, and FP16 (mixed-precision) with batch size of 2048. The 10-dimensional scVI latent representation was used for all downstream analyses.

#### Unsupervised clustering and cell type annotation

A k-nearest neighbor graph (k = 15) was constructed on the scVI latent space using rapids_singlecell (v0.12.1). UMAP visualization, Leiden clustering, and marker gene identification were performed using scanpy (v1.10.4). UMAP embeddings were generated with min_dist = 0.1, spread = 1.0, and spectral initialization. Community detection used the Leiden algorithm at resolution 0.5. Marker genes were identified via Wilcoxon rank-sum test, retaining genes with adjusted p-value < 0.05 and >30% non-zero expression in the test group to guide cell type annotation.

#### Iterative subclustering and doublet removal

Each major cell type compartment was independently subclustered using the same scVI→neighbors→UMAP→Leiden pipeline (identical hyperparameters as above). Iterative rounds of sub clustering were performed per compartment with doublets, multiplets, and contaminating populations identified by aberrant co-expression of lineage-specific markers and removed at each round.

#### Spatial niche identification

Spatial cellular niches were identified using Cell Charter (v0.3.5). A spatial neighbor graph was constructed for each TMA core using 10 nearest neighbors based on cell centroid coordinates. Neighborhood representations were computed by aggregating scVI latent embeddings over 1-hop and 2-hop spatial neighbors. The optimal number of niches was determined using Cell Charter’s ClusterAutoK module, which evaluates cluster stability across 20 independent runs for each k in the range 5–20.

#### Calculation of TMA core areas

Images were imported into QuPath for analysis using standard Hematoxylin and Eosin (H&E) stain vectors. Individual TMA cores were identified and labeled using the automated TMA dearraying tool. Parameters for core detection included a core diameter of 2.0mm, a density threshold of 5.0, and a bounds scale factor of 105.0. Following automated detection, core boundaries were manually refined where necessary. To calculate the precise tissue area within each core, a machine learning-based pixel classifier was developed. Representative regions of tissue, background, and border were manually annotated across multiple cores to create a robust training set. An artificial neural network (ANN_MLP) classifier was utilized, trained, and applied at a pixel size of 4.0μm/px. The trained classifier was applied to all detected cores. The total area for each core expressed in μm2 was automatically calculated based on the calibrated pixel size within the image metadata. Data were exported using the “Measurement Maps” feature for downstream analysis.

#### Differential cell density analysis

To test for differences in cell type, sub type and spatial niche density between treatment groups, we used edgeR’s (v4.4.0) generalized linear model (GLM) quasi-likelihood (QL) framework. Cell type counts per core were organized into a count matrix (cell types x cores). Tissue area per core (mm²) was used as the library size (equivalent to a log-area offset), allowing direct modeling of cell densities. A design matrix encoding treatment group was specified with Ctrl as the reference level. Dispersions were estimated via glmQLFit with robust = TRUE. Differential density was assessed using the QL F-test. P-values were corrected for multiple testing using the Benjamini–Hochberg (BH) method.

#### Compositional analysis

Compositional differences in cell type, sub type and spatial niche proportions between treatment groups were assessed using sccomp(v2.1.17) using the sum-constrained Beta-Binomial model. The model was specified with formula ∼ 0 + group, and the contrast groupCtrl - groupIndole was tested. Outlier cells were identified and removed using sccomp_remove_outliers. Posterior probabilities of the null hypothesis (c_pH0) and BH-corrected false discovery rates were computed for each comparison.

#### Bacterial DNA extraction

Mouse fecal pellets were collected at day 0 (before tumor cell injection), and then at days 20 and 30 after tumor cell injection. Samples were frozen on dry ice and stored at −80°C. Bacterial DNA was extracted using the QIAmp Fast DNA Stool Mini Kit (Qiagen).

#### Analysis of *Bacteroides*

To identify bacterial strain that is similar to *B. rodentium,* we performed phenotypical and genomic comparision between different *Bacteroides strains.* We analyzed genomes from BV-BRC public database (https://www.bv-brc.org/). All *Bacteroides uniformis* analysed were found to carry the TnaA enzyme (fig|411479.10.peg.1603 | BACUNI_01438 | Tryptophanase (EC 4.1.99.1), https://www.bv-brc.org/view/Feature/PATRIC.411479.10.NZ_DS362241.CDS.100381.101754.fwd and the ArAT enzyme (fig|411479.10.peg.506 | BACUNI_00099 | Aspartate/aromatic aminotransferase (EC 2.6.1.1), https://www.bv-brc.org/view/Feature/PATRIC.411479.10.NZ_DS362229.CDS.18972.20090.fwd#view_tab=overview.

#### 16S bacterial rRNA gene sequencing data processing

To determine bacterial composition, the V3/4 variable region of the 16 S rRNA gene was amplified using 16S-341F (CCTACGGGNGGCWGCAG) and 16S-785 R (GACTACHVGGGTATCTAATCC) primers.

Sequencing of pooled libraries was carried out using Illumina Nextera platform. Sequencing data were processed using Qiime2 pipeline. Raw sequencing data were processed to remove primers using cutadapt followed by denoising, merging, chimera removal to generate unique amplicon sequence variants (ASV) using the Dada2 plugin. For taxonomic assignments of these ASVs, first naive Bayes classifiers was generated on extracted V34 sequences obtained from SILVA 138.1 database using fit-classifier-naive-bayes plugin. Finally, this trained classifier was used to assign the taxonomy to the representative sequences of each ASV using feature-classifier classifiers learn plugin. Also, these representative sequences were aligned using MAFFT and phylogenetic trees both rooted and unrooted were constructed using FasTree.

Generated ASVs along with the taxonomies were used for the downstream analysis. Firstly, contaminants were identified and removed based on the prevalence of taxa in the negative control samples. Then additional filtering was done by removing ASV if it is not present in at least 10% of the samples (separately for both group; for MC the cut-off is 8 samples and for TRP the cut-off is 5 samples) followed by removing samples if they do not have sequencing depth of at least 1000 sequences. This final filtered ASV table was used for downstream statistical analysis.

Microbial community analyses were performed with the R statistical interface. Briefly, for alpha diversity, beta diversity, permutational multivariate analysis of variance (PERMANOVA), multiple R packages such as vegan, ape, phyloseq were used. For alpha diversity analysis, microbial richness was considered by identifying observed ASVs across samples and for beta diversity bray-curtis distances among samples were computed using relative abundances of each ASV in the sample. Taxonomic distribution was shown using plot bar function using phyloseq package in R. Finally, variance partition analysis was used to identify the variables which are contributing the maximum variance in the data and to detect signature taxa with maximum variance contribution. Distribution of top taxa based on the treatment were plotted separately. Kruskal–Wallis tests were used to check the statistical significance among multiple groups using the kruskalmc function of pgirmess package in R. All graphs were plotted using ggplot function in R.

#### Analysis of human databases

For analysis of *Bacteroides* and *B. uniformis* abundance in human data, we utilized microbiome abundance data from TCMA (The Cancer Microbiome Atlas)^67^. The dataset comprises microbiome abundance profiles from 620 solid tumors in TCGA studies (including HNSC, ESA, STAD, COAD, READ). The relative abundance data at both the genus level (*Bacteroides*) and the species level (*Bacteroides uniformis*) was extracted. For survival analysis, human data was updated based on harmonized outcomes reported by Liu et al.^68^ with corresponding endpoint definitions.

Kaplan-Meier survival curves were generated separately for the taxon and endpoints shown in each panel, with survival time plotted in years and p-values determined by two-sided log-rank tests. For Cox proportional hazards models, survival time was analyzed in days. The reported hazard ratios compare Present versus Absent bacterial state and were adjusted for AJCC tumor stage, age at initial pathologic diagnosis, and sex; tumor type was evaluated as an additional sensitivity covariate where applicable. Patients with missing endpoint data were excluded from the corresponding Kaplan-Meier analysis, and patients with unreported tumor stage were excluded from adjusted Cox models.

Bacterial abundance was dichotomized before survival modeling: relative abundance greater than 0 was classified as Present, and abundance equal to 0 was classified as Absent. For the genus-level Bacteroides overall survival analysis shown in Supplementary Figure 1E, TCMA genus-level relative-abundance values were collapsed to the patient level by trimming aliquot barcodes to the first 12 TCGA characters and retaining the sample with the highest abundance when multiple aliquots mapped to the same patient. In all, 503 patients had overall-survival endpoint data (Absent, n=337 with 136 deaths; Present, n=166 with 35 deaths), and 454 patients with reported tumor stage were included in the adjusted Cox model (153 deaths). This analysis included HNSC, STAD, COAD, ESCA, and READ cases.

For the species-level *B. uniformis* disease-free interval (remission) analysis shown in Figure 4D, TCMA whole-exome sequencing solid-tumor case species-level relative-abundance values were filtered to NCBI taxon ID 820 (*Bacteroides uniformis*) and merged by TCGA patient barcode. After merging clinical outcomes (per analysis performed by Liu et al.,) 224 patients had disease-free interval (remission) data (Absent, n=124 with 21 events; Present, n=100 with 9 events), and 213 patients with reported tumor stage were included in the adjusted Cox model (28 events). This species-level analysis included COAD and READ cases.

The analysis of *TnaA* abundance between responders *vs* non-responders as well as functional dysbiosis levels was performed as outlined before^69^. Briefly, data was processed using Metaphlan V4 and human V3 to obtain quantifications of Kegg Orthologs present in the microbial community. For *TnaA* quantifications, the centered log ratio abundance of the Kegg Ortholog K01667 were used. Pre-treatment samples were analyzed from the following datasets: Lee: PRJEB43119, using the PRIMM cohorts^70^, Gunjur: PRJEB49516^71^, Spencer: PRJNA770295^72^, and Routy:PRJEB22863^49^.

Analysis of tryptophan metabolism pathway activity relative to immunotherapy response was carried out in MDACC using melanoma patients data. Imputed metabolomic pathway activity for selected KEGG tryptophan metabolism enzymes^73^ derived from fecal metagenomic sequencing data. Cohort consists of melanoma patients receiving immunotherapy as previously described^72^.

#### BMDC stimulation with *B. rodentium and B. uniformis* supernatants

Bone marrow was isolated from tibiae and femurs of C57BL/6J mice and cultured 8 days in RPMI 1640 media containing 10% FBS, 1% penicillin/streptomycin, and recombinant mouse GM-CSF (20 ng/mL, BioLegend) at 37°C.

#### Detection of serum cytokines and chemokines

Serum was collected before mice were euthanized and stored at −80°C. Cytokines and chemokines were quantified using the Bio-Plex Pro Mouse Cytokine 23-plex Assay kit (BIO-RAD). All data were collected using Luminex XMAP Technology, and a MAGPIX instrument and analyzed with luminex xPONENT software.

#### Statistical analysis

Statistical analyses were performed using Prism software (version 10, GraphPad). Differences between two groups were compared using a two-tailed unpaired t-test for parametric data or the Mann-Whitney U test for non-parametric data. Tumor growth curves were analyzed using two-way ANOVA.

## Data availability

RNA sequencing data have been deposited at GEO under accession number GSE 293242.

## Acknowledgments

We thank members of the Ronai lab for extensive discussions. We also thank Lorin Chin, Mason Mandolfo and Robert Schmaltz for their assistance with bacterial cultures and animal studies. Support by the Cedars-Sinai shared resources in genomics, vivarium and microbiome studies is greatly appreciated. We gratefully acknowledge support by grant R35CA197465 (to ZAR), by gift from the Hervey Family / San Diego Foundation (to ZAR), by grant R21CA249822 (to ART and HK) and by funding from the Buffett Cancer Center funds (to ART) via the National Cancer Institute grant CA036727.

## Author contributions

X.D.O., A.R.T. and Z.A.R. conceived the study; X.D.O., K.B. and G.P. performed experiments; D.S., C.P., performed metabolite analyses; A.K.S. and A.M. analyzed bacterial 16S rRNA gene sequencing data; S.K. overseen spatial analysis,T.W.Z and A.S. performed bioinformatic analyses; NJA, JW, EEV and MPM analyzed indicated patient stool samples. E.M., S.D., A.O., M.F., O.H., A.R.T., N.J.A., G.W. and Z.A.R analyzed data; X.D.O., A.R.T., and Z.A.R. wrote the manuscript with contributions from all authors.

## Supplementary Figure Legends

**Supplementary Figure 1: *B. rodentium* colonization limited tumor growth.** Growth **(A)** and weight **(B)** of tumors in germ-free (GF) mice colonized with either ASF or ASF plus *B. rodentium, P. sartorii*, and *P. massiliensis* 14 days prior to YUMM1.5 tumor cell injection. **(C)** Weight of YUMM1.5 tumors established in GF mice colonized with either ASF or ASF plus *B. rodentium via* oral gavage 14 days prior to YUMM1.5 tumor cell injection (n = 15 mice/treatment; data represent two experiments). **(D)** Weight of YUMM1.5 tumors established in GF mice colonized with either microbiome MC608-F-a1 or MC608-F-a1 plus *B. rodentium via* oral gavage 14 days prior to YUMM1.5 tumor cell injection (n=18 mice/treatment; data represent two experiments). **(E)** Kaplan-Meier survival analysis (years) reflecting the relative abundance of the microbiome genus *Bacteroides* for overall patient survival. Biological state indicates the presence (positive) or absence (negative) of *Bacteroides* in the analysis. The log-rank test p-value is indicated in the survival plot. Cox proportional hazards model results: HR = 0.50 (95% CI: 0.33– 0.75), p = 0.001. Data were analyzed by unpaired t-test. *P < 0.05, **P < 0.005, ***P < 0.001, ****P < 0.0001 using two-tailed t-test or two-way ANOVA.

**Supplementary Figure 2: Upregulated immune signaling in mice colonized with *B. rodentium*. (A)** RT-qPCR validation of key genes functioning in anti-tumor immunity as initially identified by RNAseq. Data shown represent three analyses from each of two experiments. **(B)** RT-qPCR validation of RNAseq data of genes implicated in immune signaling (ASF, n=3; ASF plus *B. rodentium*, n=3; two independent experiments). **(C)** CD40, CD86, MHC I, and MHC II expression (MFI) on bone marrow-derived dendritic cells (BMDCs) untreated (control), stimulated with 10% of secretome from *B. rodentium,* or treated with LPS (positive control) *in vitro* for 24 hours (n=3). Data were analyzed by unpaired t-test. *P < 0.05, **P < 0.005, ***P < 0.001, by two-tailed t-test.

**Supplementary Figure 3: Metabolomic and sequencing analyses. (A)** Independent metabolomic analysis depicts the top 50 metabolic sets ranked by P-value in KEGG pathway analysis (n=6 mice/treatment). **(B)** Metabolomic analysis showing metabolites ranked by P-value (ASF, n=3; ASF plus *B. rodentium*, n=3; two independent experiments). **(C)** Independent metabolic analysis reflected in heatmap depicting clustering of metabolites enriched in indicated groups, based on KEGG pathway analysis (n=6 mice/treatment). **(D)** Effects of taurine on melanoma tumor growth based on analysis of tumor size. Taurine was provided two weeks prior to subcutaneous injection of conventional C57BL/6J mice with YUMM1.5 cells (Control, n=22; Taurine n=10). **(E)** Effects of kynurenic acid provided three days prior to injection of conventional C57BL/6J mice with tumor cells on melanoma tumor growth (Control, n=22; kynurenic acid n=12). **(F)** Abundance of the taxonomic distribution across indicated samples based on analysis of 16S rRNA bacterial gene sequencing analysis of fecal samples from the indicated groups. This analysis provided the basis for more detailed analyses of select bacterial strains that were altered in this experiment (Figure 3C). **(G)** Porcentage of relative abundance of *B. rodentium* in stool samples from WT mice administrated with normal diet or decifient tryptophan diet, at day 0 and 30 after tumor inoculation (n=5 mice per treatment). Data were analyzed by unpaired t-test. *P < 0.05 **P < 0.005, ***P < 0.001, ****P < 0.0001 by two-tailed t-test or two-way ANOVA.

**Supplementary Figure 4: *B. uniformis* colonization limited tumor growth. (A)** Weight of YUMM1.5 tumors in GF mice colonized with either microbiome MC608-F-a1 or MC608-F-a1 plus *B. uniformis*. **(B)** Weight of YUMM 1.5 tumors in conventional C57BL/6J mice colonized with either *B. rodentium* or *B. uniformis*. **(C)** RT-qPCR analysis validating key genes identified by RNAseq in GF mice colonized with MC608-F-a1, MC608-F-a1 plus *B. rodentium* or MC608-F-a1 plus *B. uniformis* (n=4 mice/treatment). **(D)** RT-qPCR analysis of genes identified by RNAseq as implicated in control of immune signaling in GF mice colonized with MC608-F-a1, MC608-F-a1 plus *B. rodentium*, or MC608-F-a1 plus *B. uniformis* (n=4/treatment). **(E)** CD40, CD86, MHC I, and MHC II expression (MFI) on bone marrow-derived dendritic cells (BMDCs) untreated (control), stimulated with 10% of secretome from *B. uniformis*, or treated with LPS (positive control) *in vitro* for 24 hours (n=3). Data shown represent two experiments. Data were analyzed by unpaired t-test. *P < 0.05 **P < 0.005, ***P < 0.001, ****P < 0.0001 by two-tailed t-test.

**Supplementary Figure 5: *B. uniformis* colonization limited tumor growth requires the tryptophanase (*TnaA*) gene. (A)** PCR validation of mutant tryptophanase in *B. uniformis,* and illustration of the tryptophanase gene deletion by allelic exchange **(B)** Validation of mutant tryptophanase activity in *B. uniformis* was assessed *in vitro* in high- or low-tryptophan media. **(C)** Weight of YUMM 1.5 tumors in GF mice that were colonized with either ASF, ASF plus *B. uniformis,* or ASF plus *B. uniformis* tryptophanase (*TnaA*) mutant. **(D)** Weight of YUMM 1.5 tumors in GF mice that were colonized with either *B. uniformis* or *B. uniformis TnaA* mutant. **(E)** Quantification of indicated indoles metabolites in cecal samples from GF mice colonized with *B. uniformis* or *B. uniformis tryptophanase* mutant (n = 5 mice/treatment). **(F)** Serum cytokines levels in GF mice colonized with ASF, ASF plus *B. rodentium* (n=5 mice/treatment) or ASF, ASF plus *B. uniformis,* ASF plus *B. uniformis TnaA* mutant (n=5 mice/treatment); conventional C57BL/6J mice, colonized with either *B. rodentium* or *B. uniformis* (n=4 mice/treatment). **(G)** Weight of YUMM 1.5 tumors in conventional C57BL/6J mice that were treated with *B. uniformis,* kynurenine, indoxyl sulfate, indole pyruvate, indole lactic acid or indole. **(H)** RT-qPCR of genes implicated in immune signaling (Control, n=3; Indoles, n=3; two independent experiments). **(I)** Quantification of tumor infiltration of CD8^+^ and CD4^+^ T cells, CD44^+^, CD69^+^, CD62L^+^, TIM 3^+^, PD1^+^, GZMB^+^, TNFα^+^ and IFNγ^+^ on CD4^+^ and CD8^+^ T cell 12 days after injection of YUMM1.5 cells into conventional C57BL/6J mice treated with indoles (daily *via* oral gavage starting one day after tumor inoculation until end of the experiment), (n =8 mice/treatment). Data were analyzed by unpaired t-test. *P < 0.05, **P < 0.005, ***P < 0.001, ****P < 0.0001 by two-tailed t-test or two-way ANOVA.

**Supplementary Figure 6:** Inhibition of tumor growth by *B. rodentium* or indole administration appears to be AhR, PXR and GPCR independent. (A) RT-qPCR of genes implicated in AhR signaling (ASF, n=6; ASF plus *B. rodentium,* n=6). **(B)** RT-qPCR of genes implicated in AhR signaling (Control, n=5; Indoles, n=5). **(C)** Tumor growth in GF mice colonized with either ASF, ASF plus *B. rodentium,* 14 days prior to YUMM1.5 tumor cell injection, treated with AhR agonist (TCDD) or vehicle. (n = 15 mice/treatment). **(D)** RT-qPCR of genes implicated in GPCR signaling (Control, n=5; Indoles, n=5) **(E)** RT-qPCR of genes implicated in PXR signaling (Control, n=5; Indoles, n=5). Data were analyzed by unpaired t-test. *P < 0.05, **P < 0.005, ***P < 0.001, ****P < 0.0001 by two-tailed t-test or two-way ANOVA.

## Supplemental Tables

**Supplemental table 1:** List of primers used for qRT-PCR analysis.

**Supplemental table 2:** Standards used for Gas Chromatography-Mass Spectrometry (GC-MS) Analysis.

**Supplemental table 3:** List of different metabolites analyzed from cecal samples from mice colonized with ASF or ASF plus *B. rodentium* by Gas Chromatography-Mass Spectrometry (GC-MS). The table include in the column the index (A unique identifier for each detected compound in the dataset); compounds (name of the metabolite assigned to that GC-Ms peak); Class I (A broad chemical category); Class II (A more specific subclass of Class I); Q1(Da) (The mass-to-charge ratio (m/z) of the ion selected in the first quadrupole (Q1)); Molecular weight (Da) (The theoretical molecular mass ot the compound); Ionizacion model (How the molecule was ionized by the mass sprectrometer); Formula (Chemcal formula of the compound) and level (The confidence level of the compound identification) for each of the metabolites analyzed. Green highlight provide measures of the metabolites associated to ASF group, while the orange highlight provide measures of the metabolites associated with ASF plus *B. rodentium* group. (6 mice per group).

**Supplemental Table 4:** Binary phenotype matrix comparison of *B. rodentium* and other *Bacteroides* strains. The table compares the phenotypic metabolites of each of the *Bacteroides* strains, with *B. rodentium.* The metabolites identified are highlighted in green (1) whereas orange highlights the metabolites that are absent (0). Intermediate values (e.g 0.7) present partial or weak expression.

**Supplemental Table 5:** List of analyzed genomes from BV-BRC public database (https://www.bv-brc.org/). The table include ID (The unique genome identifier in BV-BRC public database); Genus (The genus name of the organism); Species (The species name); and mc-which present the name of the bacterial strain; Similarity is presented by normalized value reflecting degree of similarity with *B. rodentium* genome.

## References

1. Arozarena, I., and Wellbrock, C. (2017). Overcoming resistance to BRAF inhibitors. Ann Transl Med 5, 387. 10.21037/atm.2017.06.09.

2. Long, G.V., Carlino, M.S., Au-Yeung, G., Spillane, A.J., Shannon, K.F., Gyorki, D.E., Hsiao, E., Kapoor, R., Thompson, J.R., Batula, I., et al. (2024). Neoadjuvant pembrolizumab, dabrafenib and trametinib in BRAF(V600)-mutant resectable melanoma: the randomized phase 2 NeoTrio trial. Nat Med 30, 2540–2548. 10.1038/s41591-024-03077-5.

3. Routy, B., Jackson, T., Mahlmann, L., Baumgartner, C.K., Blaser, M., Byrd, A., Corvaia, N., Couts, K., Davar, D., Derosa, L., et al. (2024). Melanoma and microbiota: Current understanding and future directions. Cancer Cell 42, 16–34. 10.1016/j.ccell.2023.12.003.

4. Li, Y., Tinoco, R., Elmen, L., Segota, I., Xian, Y., Fujita, Y., Sahu, A., Zarecki, R., Marie, K., Feng, Y., et al. (2019). Gut microbiota dependent anti-tumor immunity restricts melanoma growth in Rnf5(-/-) mice. Nat Commun 10, 1492. 10.1038/s41467-019-09525-y.

5. Gopalakrishnan, V., Spencer, C.N., Nezi, L., Reuben, A., Andrews, M.C., Karpinets, T.V., Prieto, P.A., Vicente, D., Hoffman, K., Wei, S.C., et al. (2018). Gut microbiome modulates response to anti-PD-1 immunotherapy in melanoma patients. Science 359, 97–103. 10.1126/science.aan4236.

6. Tanoue, T., Morita, S., Plichta, D.R., Skelly, A.N., Suda, W., Sugiura, Y., Narushima, S., Vlamakis, H., Motoo, I., Sugita, K., et al. (2019). A defined commensal consortium elicits CD8 T cells and anti-cancer immunity. Nature 565, 600–605. 10.1038/s41586-019-0878-z.

7. Sivan, A., Corrales, L., Hubert, N., Williams, J.B., Aquino-Michaels, K., Earley, Z.M., Benyamin, F.W., Lei, Y.M., Jabri, B., Alegre, M.L., et al. (2015). Commensal Bifidobacterium promotes antitumor immunity and facilitates anti-PD-L1 efficacy. Science 350, 1084–1089. 10.1126/science.aac4255.

8. Zagato, E., Pozzi, C., Bertocchi, A., Schioppa, T., Saccheri, F., Guglietta, S., Fosso, B., Melocchi, L., Nizzoli, G., Troisi, J., et al. (2020). Endogenous murine microbiota member Faecalibaculum rodentium and its human homologue protect from intestinal tumour growth. Nat Microbiol 5, 511–524. 10.1038/s41564-019-0649-5.

9. Kuang, E., Qi, J., and Ronai, Z. (2013). Emerging roles of E3 ubiquitin ligases in autophagy. Trends Biochem Sci 38, 453–460. 10.1016/j.tibs.2013.06.008.

10. Tcherpakov, M., Delaunay, A., Toth, J., Kadoya, T., Petroski, M.D., and Ronai, Z.A. (2009). Regulation of endoplasmic reticulum-associated degradation by RNF5-dependent ubiquitination of JNK-associated membrane protein (JAMP). J Biol Chem 284, 12099–12109. 10.1074/jbc.M808222200.

11. Fujita, Y., Khateb, A., Li, Y., Tinoco, R., Zhang, T., Bar-Yoseph, H., Tam, M.A., Chowers, Y., Sabo, E., Gerassy-Vainberg, S., et al. (2018). Regulation of S100A8 Stability by RNF5 in Intestinal Epithelial Cells Determines Intestinal Inflammation and Severity of Colitis. Cell Rep 24, 3296–3311 e3296. 10.1016/j.celrep.2018.08.057.

12. Jeon, Y.J., Khelifa, S., Ratnikov, B., Scott, D.A., Feng, Y., Parisi, F., Ruller, C., Lau, E., Kim, H., Brill, L.M., et al. (2015). Regulation of glutamine carrier proteins by RNF5 determines breast cancer response to ER stress-inducing chemotherapies. Cancer Cell 27, 354–369. 10.1016/j.ccell.2015.02.006.

13. Yan, J., Chen, D., Ye, Z., Zhu, X., Li, X., Jiao, H., Duan, M., Zhang, C., Cheng, J., Xu, L., et al. (2024). Molecular mechanisms and therapeutic significance of Tryptophan Metabolism and signaling in cancer. Mol Cancer 23, 241. 10.1186/s12943-024-02164-y.

14. Su, X., Gao, Y., and Yang, R. (2022). Gut Microbiota-Derived Tryptophan Metabolites Maintain Gut and Systemic Homeostasis. Cells 11. 10.3390/cells11152296.

15. Gao, J., Xu, K., Liu, H., Liu, G., Bai, M., Peng, C., Li, T., and Yin, Y. (2018). Impact of the Gut Microbiota on Intestinal Immunity Mediated by Tryptophan Metabolism. Front Cell Infect Microbiol 8, 13. 10.3389/fcimb.2018.00013.

16. Baruch, E.N., Youngster, I., Ben-Betzalel, G., Ortenberg, R., Lahat, A., Katz, L., Adler, K., Dick-Necula, D., Raskin, S., Bloch, N., et al. (2021). Fecal microbiota transplant promotes response in immunotherapy-refractory melanoma patients. Science 371, 602–609. 10.1126/science.abb5920.

17. Andrews, M.C., Duong, C.P.M., Gopalakrishnan, V., Iebba, V., Chen, W.S., Derosa, L., Khan, M.A.W., Cogdill, A.P., White, M.G., Wong, M.C., et al. (2021). Gut microbiota signatures are associated with toxicity to combined CTLA-4 and PD-1 blockade. Nat Med 27, 1432–1441. 10.1038/s41591-021-01406-6.

18. Baruch, E.N., Wang, J., and Wargo, J.A. (2021). Gut Microbiota and Antitumor Immunity: Potential Mechanisms for Clinical Effect. Cancer Immunol Res 9, 365–370. 10.1158/2326-6066.CIR-20-0877.

19. Hezaveh, K., Shinde, R.S., Klotgen, A., Halaby, M.J., Lamorte, S., Ciudad, M.T., Quevedo, R., Neufeld, L., Liu, Z.Q., Jin, R., et al. (2022). Tryptophan-derived microbial metabolites activate the aryl hydrocarbon receptor in tumor-associated macrophages to suppress anti-tumor immunity. Immunity 55, 324–340 e328. 10.1016/j.immuni.2022.01.006.

20. Kim, M., and Tomek, P. (2021). Tryptophan: A Rheostat of Cancer Immune Escape Mediated by Immunosuppressive Enzymes IDO1 and TDO. Front Immunol 12, 636081. 10.3389/fimmu.2021.636081.

21. Mondanelli, G., Ugel, S., Grohmann, U., and Bronte, V. (2017). The immune regulation in cancer by the amino acid metabolizing enzymes ARG and IDO. Curr Opin Pharmacol 35, 30–39. 10.1016/j.coph.2017.05.002.

22. Segura Munoz, R.R., Mantz, S., Martinez, I., Li, F., Schmaltz, R.J., Pudlo, N.A., Urs, K., Martens, E.C., Walter, J., and Ramer-Tait, A.E. (2022). Experimental evaluation of ecological principles to understand and modulate the outcome of bacterial strain competition in gut microbiomes. ISME J 16, 1594–1604. 10.1038/s41396-022-01208-9.

23. Lagkouvardos, I., Pukall, R., Abt, B., Foesel, B.U., Meier-Kolthoff, J.P., Kumar, N., Bresciani, A., Martinez, I., Just, S., Ziegler, C., et al. (2016). The Mouse Intestinal Bacterial Collection (miBC) provides host-specific insight into cultured diversity and functional potential of the gut microbiota. Nat Microbiol 1, 16131. 10.1038/nmicrobiol.2016.131.

24. Martinez, I., Maldonado-Gomez, M.X., Gomes-Neto, J.C., Kittana, H., Ding, H., Schmaltz, R., Joglekar, P., Cardona, R.J., Marsteller, N.L., Kembel, S.W., et al. (2018). Experimental evaluation of the importance of colonization history in early-life gut microbiota assembly. Elife 7. 10.7554/eLife.36521.

25. Rad Pour, S., Morikawa, H., Kiani, N.A., Yang, M., Azimi, A., Shafi, G., Shang, M., Baumgartner, R., Ketelhuth, D.F.J., Kamleh, M.A., et al. (2019). Exhaustion of CD4+ T-cells mediated by the Kynurenine Pathway in Melanoma. Sci Rep 9, 12150. 10.1038/s41598-019-48635-x.

26. Jiang, Y., Tao, Q., Qiao, X., Yang, Y., Peng, C., Han, M., Dong, K., Zhang, W., Xu, M., Wang, D., et al. (2025). Targeting amino acid metabolism to inhibit gastric cancer progression and promote anti-tumor immunity: a review. Front Immunol 16, 1508730. 10.3389/fimmu.2025.1508730.

27. Li, L., and McAllister, F. (2022). Too much water drowned the miller: Akkermansia determines immunotherapy responses. Cell Rep Med 3, 100642. 10.1016/j.xcrm.2022.100642.

28. Hou, Y., Li, J., and Ying, S. (2023). Tryptophan Metabolism and Gut Microbiota: A Novel Regulatory Axis Integrating the Microbiome, Immunity, and Cancer. Metabolites 13. 10.3390/metabo13111166.

29. Jin, U.H., Lee, S.O., Sridharan, G., Lee, K., Davidson, L.A., Jayaraman, A., Chapkin, R.S., Alaniz, R., and Safe, S. (2014). Microbiome-derived tryptophan metabolites and their aryl hydrocarbon receptor-dependent agonist and antagonist activities. Mol Pharmacol 85, 777–788. 10.1124/mol.113.091165.

30. Karagiannidis, I., Salataj, E., Said Abu Egal, E., and Beswick, E.J. (2021). G-CSF in tumors: Aggressiveness, tumor microenvironment and immune cell regulation. Cytokine 142, 155479. 10.1016/j.cyto.2021.155479.

31. Mouchemore, K.A., and Anderson, R.L. (2021). Immunomodulatory effects of G-CSF in cancer: Therapeutic implications. Semin Immunol 54, 101512. 10.1016/j.smim.2021.101512.

32. Son, D.S., Parl, A.K., Rice, V.M., and Khabele, D. (2007). Keratinocyte chemoattractant (KC)/human growth-regulated oncogene (GRO) chemokines and pro-inflammatory chemokine networks in mouse and human ovarian epithelial cancer cells. Cancer Biol Ther 6, 1302–1312. 10.4161/cbt.6.8.4506.

33. Korbecki, J., Bosiacki, M., Barczak, K., Lagocka, R., Chlubek, D., and Baranowska-Bosiacka, I. (2023). The Clinical Significance and Role of CXCL1 Chemokine in Gastrointestinal Cancers. Cells 12 10.3390/cells12101406.

34. Elias, A.E., McBain, A.J., Aldehalan, F.A., Taylor, G., and O’Neill, C.A. (2024). Activation of the aryl hydrocarbon receptor via indole derivatives is a common feature in skin bacterial isolates. J Appl Microbiol 135 10.1093/jambio/lxae273.

35. Hubbard, T.D., Murray, I.A., Bisson, W.H., Lahoti, T.S., Gowda, K., Amin, S.G., Patterson, A.D., and Perdew, G.H. (2015). Adaptation of the human aryl hydrocarbon receptor to sense microbiota-derived indoles. Sci Rep 5, 12689. 10.1038/srep12689.

36. Svetocheva, Z.F., and Krasnikov, P.G. (1985). [Clinical characteristics of inflammatory diseases of the optic nerve in children]. Oftalmol Zh, 327–329.

37. Bender, M.J., McPherson, A.C., Phelps, C.M., Pandey, S.P., Laughlin, C.R., Shapira, J.H., Medina Sanchez, L., Rana, M., Richie, T.G., Mims, T.S., et al. (2023). Dietary tryptophan metabolite released by intratumoral Lactobacillus reuteri facilitates immune checkpoint inhibitor treatment. Cell 186, 1846–1862 e1826. 10.1016/j.cell.2023.03.011.

38. Kocher, F., Amann, A., Zimmer, K., Geisler, S., Fuchs, D., Pichler, R., Wolf, D., Kurz, K., Seeber, A., and Pircher, A. (2021). High indoleamine-2,3-dioxygenase 1 (IDO) activity is linked to primary resistance to immunotherapy in non-small cell lung cancer (NSCLC). Transl Lung Cancer Res 10, 304–313. 10.21037/tlcr-20-380.

39. Chen, X.X., Ju, Q., Qiu, D., Zhou, Y., Wang, Y., Zhang, X.X., Li, J.G., Wang, M., Chang, N., Xu, X.R., et al. (2025). Microbial dysbiosis with tryptophan metabolites alteration in lower respiratory tract is associated with clinical responses to anti-PD-1 immunotherapy in advanced non-small cell lung cancer. Cancer Immunol Immunother 74, 140. 10.1007/s00262-025-03996-3.

40. Hu, Y., Xu, X., Zhong, H., Ding, C., Zhang, S., Qin, W., Zhang, E., Shu, D., Yu, M., Naijipu, A., et al. (2025). Integrated single cell and bulk RNA sequencing analyses reveal the impact of tryptophan metabolism on prognosis and immunotherapy in colon cancer. Sci Rep 15, 12496. 10.1038/s41598-025-85893-4.

41. Zitvogel, L., Ayyoub, M., Routy, B., and Kroemer, G. (2016). Microbiome and Anticancer Immunosurveillance. Cell 165, 276–287. 10.1016/j.cell.2016.03.001.

42. Terrisse, S., Zitvogel, L., and Kroemer, G. (2022). Effects of the intestinal microbiota on prostate cancer treatment by androgen deprivation therapy. Microb Cell 9, 202–206. 10.15698/mic2022.12.787.

43. Sepich-Poore, G.D., Zitvogel, L., Straussman, R., Hasty, J., Wargo, J.A., and Knight, R. (2021). The microbiome and human cancer. Science 371. 10.1126/science.abc4552.

44. Pitt, J.M., Vetizou, M., Waldschmitt, N., Kroemer, G., Chamaillard, M., Boneca, I.G., and Zitvogel, L. (2016). Fine-Tuning Cancer Immunotherapy: Optimizing the Gut Microbiome. Cancer Res 76, 4602–4607. 10.1158/0008-5472.CAN-16-0448.

45. Tintelnot, J., Xu, Y., Lesker, T.R., Schonlein, M., Konczalla, L., Giannou, A.D., Pelczar, P., Kylies, D., Puelles, V.G., Bielecka, A.A., et al. (2023). Microbiota-derived 3-IAA influences chemotherapy efficacy in pancreatic cancer. Nature 615, 168–174. 10.1038/s41586-023-05728-y.

46. Sugimura, N., Li, Q., Chu, E.S.H., Lau, H.C.H., Fong, W., Liu, W., Liang, C., Nakatsu, G., Su, A.C.Y., Coker, O.O., et al. (2021). Lactobacillus gallinarum modulates the gut microbiota and produces anti-cancer metabolites to protect against colorectal tumourigenesis. Gut 71, 2011–2021. 10.1136/gutjnl-2020-323951.

47. Pietrzak, B., Tomela, K., Olejnik-Schmidt, A., Galus, L., Mackiewicz, J., Kaczmarek, M., Mackiewicz, A., and Schmidt, M. (2022). A Clinical Outcome of the Anti-PD-1 Therapy of Melanoma in Polish Patients Is Mediated by Population-Specific Gut Microbiome Composition. Cancers (Basel) 14. 10.3390/cancers14215369.

48. Shen, Z., Gu, X., Cao, H., Mao, W., Yang, L., He, M., Zhang, R., Zhou, Y., Liu, K., Wang, L., et al. (2021). Characterization of microbiota in acute leukemia patients following successful remission induction chemotherapy without antimicrobial prophylaxis. Int Microbiol 24, 263–273. 10.1007/s10123-021-00163-3.

49. Routy, B., Le Chatelier, E., Derosa, L., Duong, C.P.M., Alou, M.T., Daillere, R., Fluckiger, A., Messaoudene, M., Rauber, C., Roberti, M.P., et al. (2018). Gut microbiome influences efficacy of PD-1-based immunotherapy against epithelial tumors. Science 359, 91–97. 10.1126/science.aan3706.

50. Agusti, A., Campillo, I., Balzano, T., Benitez-Paez, A., Lopez-Almela, I., Romani-Perez, M., Forteza, J., Felipo, V., Avena, N.M., and Sanz, Y. (2021). Bacteroides uniformis CECT 7771 Modulates the Brain Reward Response to Reduce Binge Eating and Anxiety-Like Behavior in Rat. Mol Neurobiol 58, 4959–4979. 10.1007/s12035-021-02462-2.

51. Ben-Shaanan, T.L., Schiller, M., Azulay-Debby, H., Korin, B., Boshnak, N., Koren, T., Krot, M., Shakya, J., Rahat, M.A., Hakim, F., and Rolls, A. (2018). Modulation of anti-tumor immunity by the brain’s reward system. Nat Commun 9, 2723. 10.1038/s41467-018-05283-5.

52. Zhang, H., Hong, Y., Wu, T., Ben, E., Li, S., Hu, L., and Xie, T. (2024). Role of gut microbiota in regulating immune checkpoint inhibitor therapy for glioblastoma. Front Immunol 15, 1401967. 10.3389/fimmu.2024.1401967.

53. Yan, Y., Lei, Y., Qu, Y., Fan, Z., Zhang, T., Xu, Y., Du, Q., Brugger, D., Chen, Y., Zhang, K., and Zhang, E. (2023). Bacteroides uniformis-induced perturbations in colonic microbiota and bile acid levels inhibit TH17 differentiation and ameliorate colitis developments. NPJ Biofilms Microbiomes 9, 56. 10.1038/s41522-023-00420-5.

54. Sharpe, A.H., and Pauken, K.E. (2018). The diverse functions of the PD1 inhibitory pathway. Nat Rev Immunol 18, 153–167. 10.1038/nri.2017.108.

55. Paluch, C., Santos, A.M., Anzilotti, C., Cornall, R.J., and Davis, S.J. (2018). Immune Checkpoints as Therapeutic Targets in Autoimmunity. Front Immunol 9, 2306. 10.3389/fimmu.2018.02306.

56. Gomes-Neto, J.C., Mantz, S., Held, K., Sinha, R., Segura Munoz, R.R., Schmaltz, R., Benson, A.K., Walter, J., and Ramer-Tait, A.E. (2017). A real-time PCR assay for accurate quantification of the individual members of the Altered Schaedler Flora microbiota in gnotobiotic mice. J Microbiol Methods 135, 52–62. 10.1016/j.mimet.2017.02.003.

57. Meeth, K., Wang, J.X., Micevic, G., Damsky, W., and Bosenberg, M.W. (2016). The YUMM lines: a series of congenic mouse melanoma cell lines with defined genetic alterations. Pigment Cell Melanoma Res 29, 590–597. 10.1111/pcmr.12498.

58. Bindels, L.B., Segura Munoz, R.R., Gomes-Neto, J.C., Mutemberezi, V., Martinez, I., Salazar, N., Cody, E.A., Quintero-Villegas, M.I., Kittana, H., de Los Reyes-Gavilan, C.G., et al. (2017). Resistant starch can improve insulin sensitivity independently of the gut microbiota. Microbiome 5, 12. 10.1186/s40168-017-0230-5.

59. Braun, L.M., Giesler, S., Andrieux, G., Riemer, R., Talvard-Balland, N., Duquesne, S., Ruckert, T., Unger, S., Kreutmair, S., Zwick, M., et al. (2025). Adiponectin reduces immune checkpoint inhibitor-induced inflammation without blocking anti-tumor immunity. Cancer Cell 43, 269–291 e219. 10.1016/j.ccell.2025.01.004.

60. Scott, D.A. (2021). Analysis of Melanoma Cell Glutamine Metabolism by Stable Isotope Tracing and Gas Chromatography-Mass Spectrometry. Methods Mol Biol 2265, 91–110. 10.1007/978-1-0716-1205-7_7.

61. Liao, Y., Smyth, G.K., and Shi, W. (2014). featureCounts: an efficient general purpose program for assigning sequence reads to genomic features. Bioinformatics 30, 923–930. 10.1093/bioinformatics/btt656.

62. Dobin, A., Davis, C.A., Schlesinger, F., Drenkow, J., Zaleski, C., Jha, S., Batut, P., Chaisson, M., and Gingeras, T.R. (2013). STAR: ultrafast universal RNA-seq aligner. Bioinformatics 29, 15–21. 10.1093/bioinformatics/bts635.

63. Li, B., and Dewey, C.N. (2011). RSEM: accurate transcript quantification from RNA-Seq data with or without a reference genome. BMC Bioinformatics 12, 323. 10.1186/1471-2105-12-323.

64. Ewels, P., Magnusson, M., Lundin, S., and Kaller, M. (2016). MultiQC: summarize analysis results for multiple tools and samples in a single report. Bioinformatics 32, 3047–3048. 10.1093/bioinformatics/btw354.

65. Li, X., Cooper, N.G.F., O’Toole, T.E., and Rouchka, E.C. (2020). Choice of library size normalization and statistical methods for differential gene expression analysis in balanced two-group comparisons for RNA-seq studies. BMC Genomics 21, 75. 10.1186/s12864-020-6502-7.

66. Subramanian, A., Tamayo, P., Mootha, V.K., Mukherjee, S., Ebert, B.L., Gillette, M.A., Paulovich, A., Pomeroy, S.L., Golub, T.R., Lander, E.S., and Mesirov, J.P. (2005). Gene set enrichment analysis: a knowledge-based approach for interpreting genome-wide expression profiles. Proc Natl Acad Sci U S A 102, 15545–15550. 10.1073/pnas.0506580102.

67. Dohlman, A.B., Arguijo Mendoza, D., Ding, S., Gao, M., Dressman, H., Iliev, I.D., Lipkin, S.M., and Shen, X. (2021). The cancer microbiome atlas: a pan-cancer comparative analysis to distinguish tissue-resident microbiota from contaminants. Cell Host Microbe 29, 281–298 e285. 10.1016/j.chom.2020.12.001.

68. Liu, J., Lichtenberg, T., Hoadley, K.A., Poisson, L.M., Lazar, A.J., Cherniack, A.D., Kovatich, A.J., Benz, C.C., Levine, D.A., Lee, A.V., et al. (2018). An Integrated TCGA Pan-Cancer Clinical Data Resource to Drive High-Quality Survival Outcome Analytics. Cell 173, 400–416 e411. 10.1016/j.cell.2018.02.052.

69. Mimpen, I.L., Battaglia, T.W., Parra-Martinez, M., Toner-Bartelds, C., Zeverijn, L.J., Geurts, B.S., Verkerk, K., Hoes, L.R., van Renterghem, A.W.J., Noe, M., et al. (2026). Microbial Metabolic Pathways Guide Response to Immune Checkpoint Blockade Therapy. Cancer Discov 16, 95–113. 10.1158/2159-8290.CD-24-1669.

70. Lee, K.A., Thomas, A.M., Bolte, L.A., Bjork, J.R., de Ruijter, L.K., Armanini, F., Asnicar, F., Blanco-Miguez, A., Board, R., Calbet-Llopart, N., et al. (2022). Cross-cohort gut microbiome associations with immune checkpoint inhibitor response in advanced melanoma. Nat Med 28, 535–544. 10.1038/s41591-022-01695-5.

71. Gunjur, A., Shao, Y., Rozday, T., Klein, O., Mu, A., Haak, B.W., Markman, B., Kee, D., Carlino, M.S., Underhill, C., et al. (2024). A gut microbial signature for combination immune checkpoint blockade across cancer types. Nat Med 30, 797–809. 10.1038/s41591-024-02823-z.

72. Spencer, C.N., McQuade, J.L., Gopalakrishnan, V., McCulloch, J.A., Vetizou, M., Cogdill, A.P., Khan, M.A.W., Zhang, X., White, M.G., Peterson, C.B., et al. (2021). Dietary fiber and probiotics influence the gut microbiome and melanoma immunotherapy response. Science 374, 1632–1640. 10.1126/science.aaz7015.

73. Liu, Z.Q., Ciudad, M.T., and McGaha, T.L. (2025). New insights into tryptophan metabolism in cancer. Trends Cancer 11, 629–641. 10.1016/j.trecan.2025.03.008.

