## Supplementary material for "Tryptophan degradation by intestinal Bacteroides induces anti-tumor immunity and limits melanoma growth": RONAI SUPP FIG

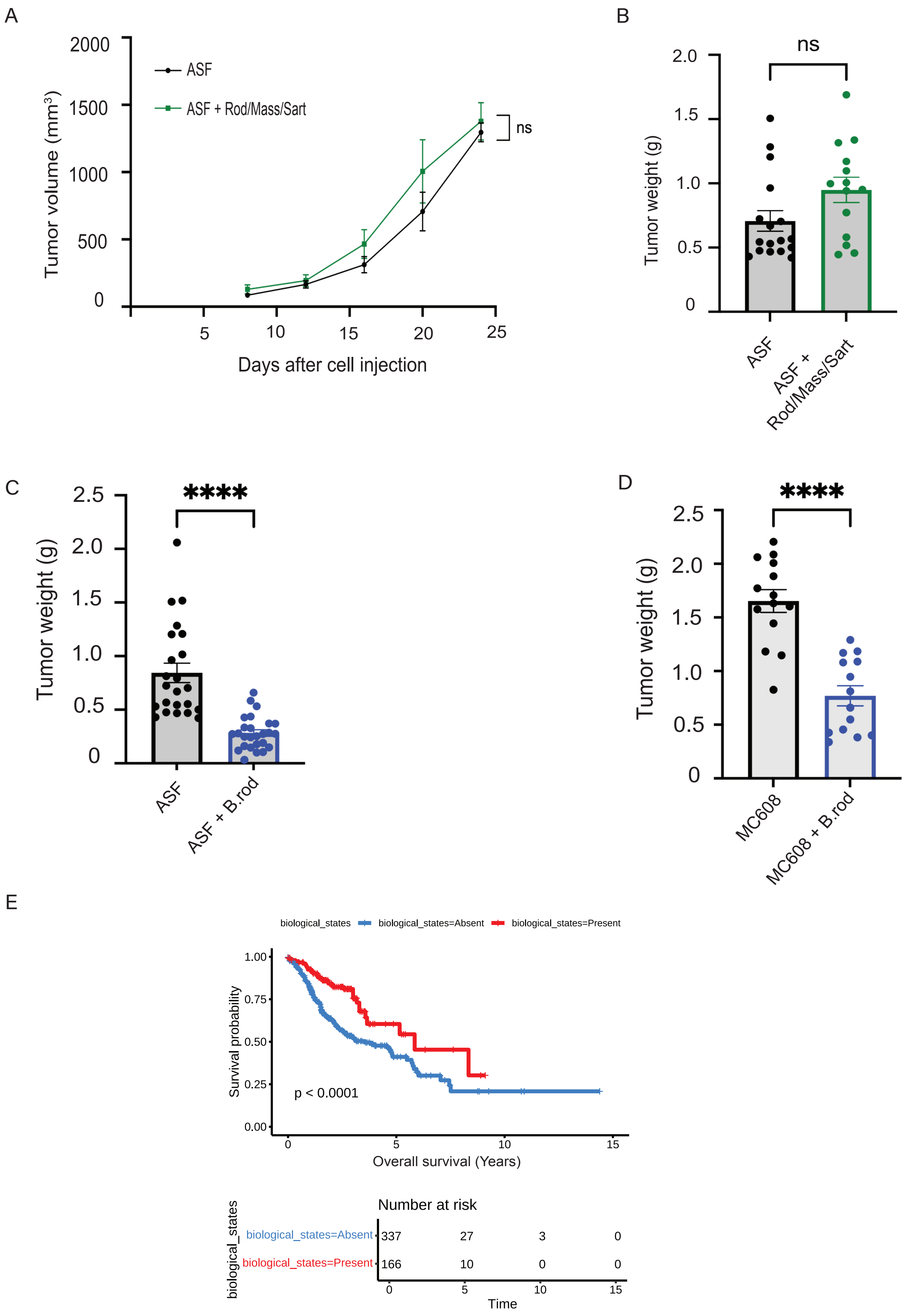


Diaz Olea et al., Supp Fig 1

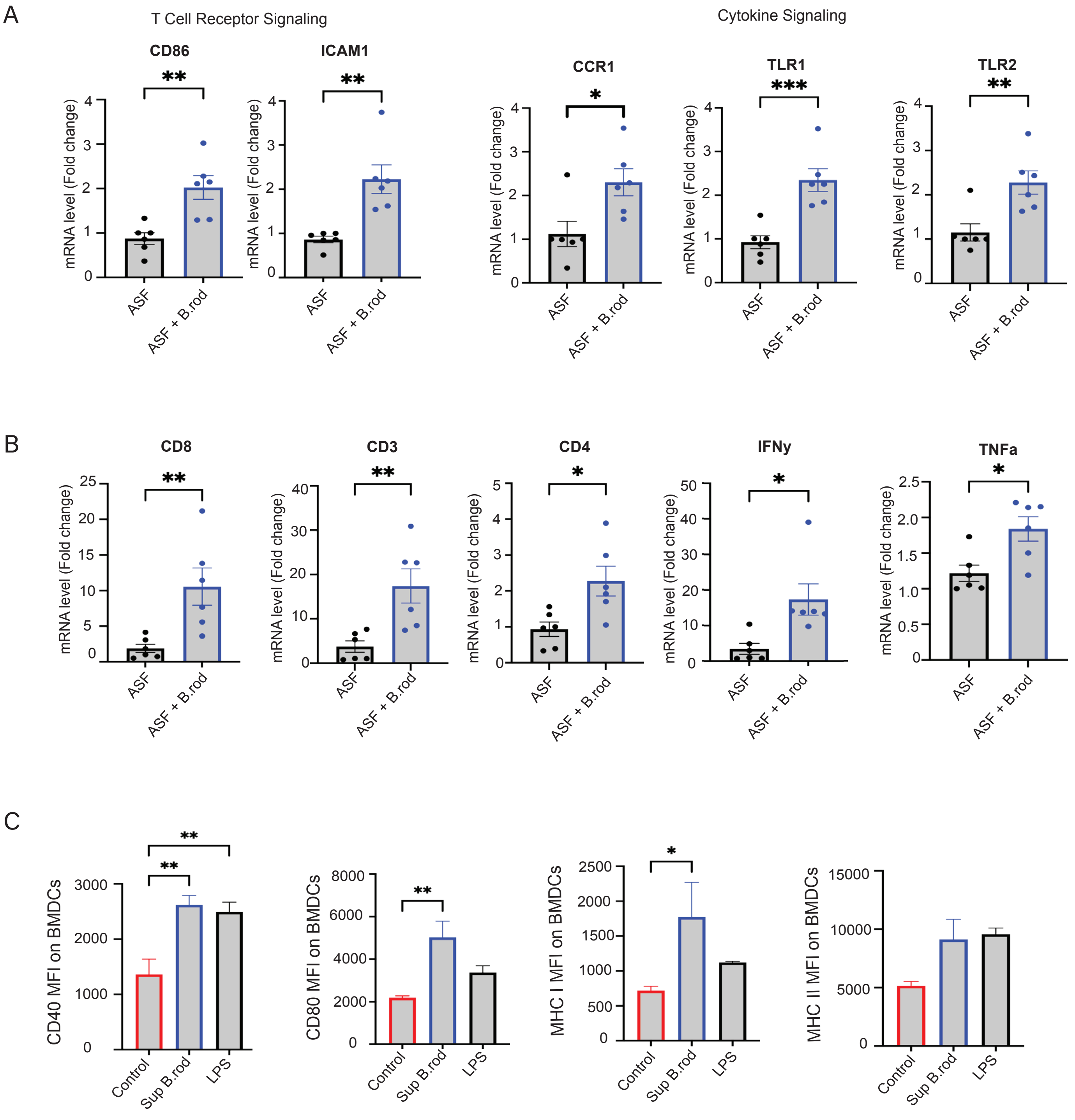


Diaz Olea et al., Supp Fig 2

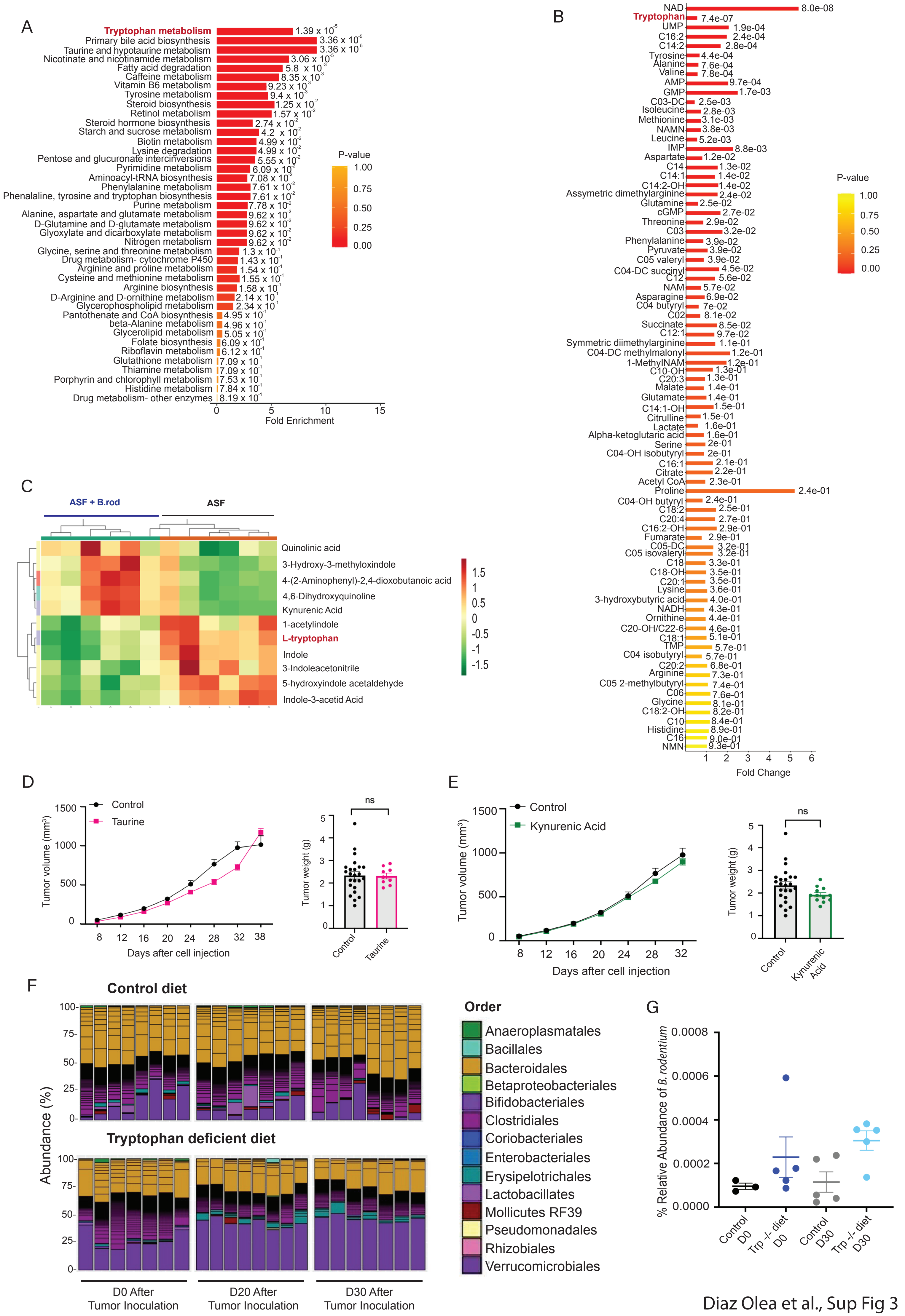


Diaz Olea et al., Sup Fig 3

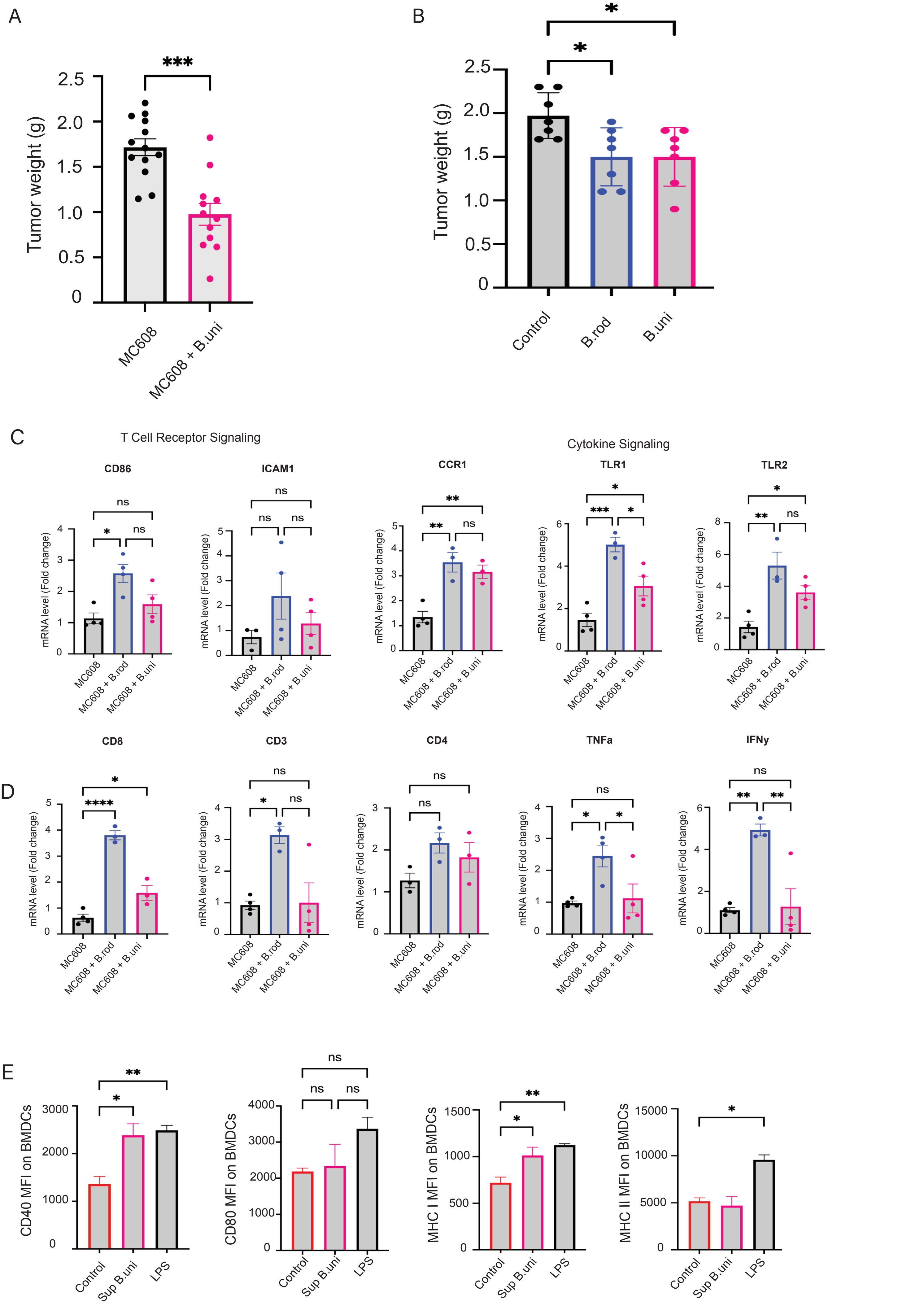


Diaz Olea et al., Sup Fig 4

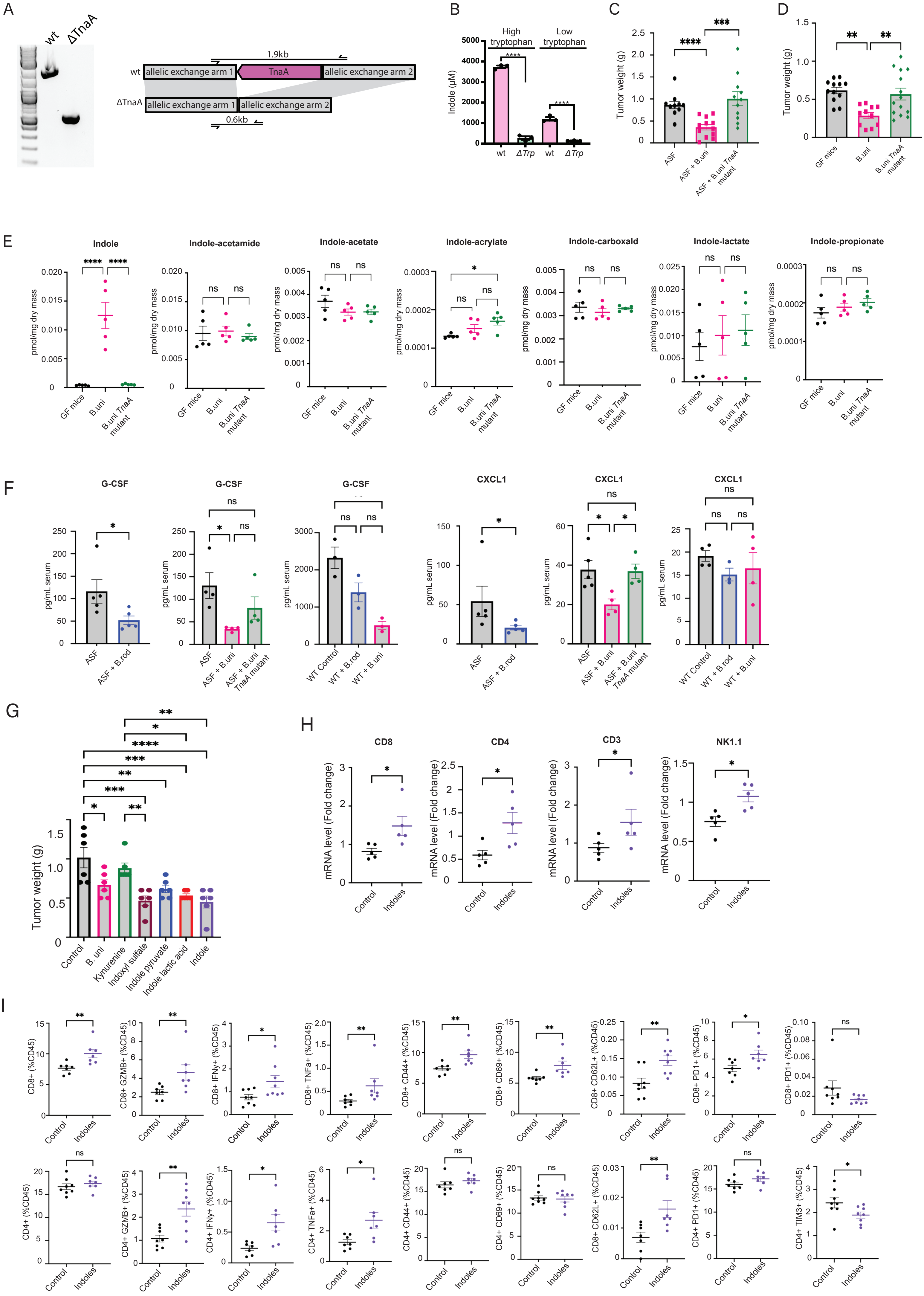


Diaz Olea et al., Sup Fig 5

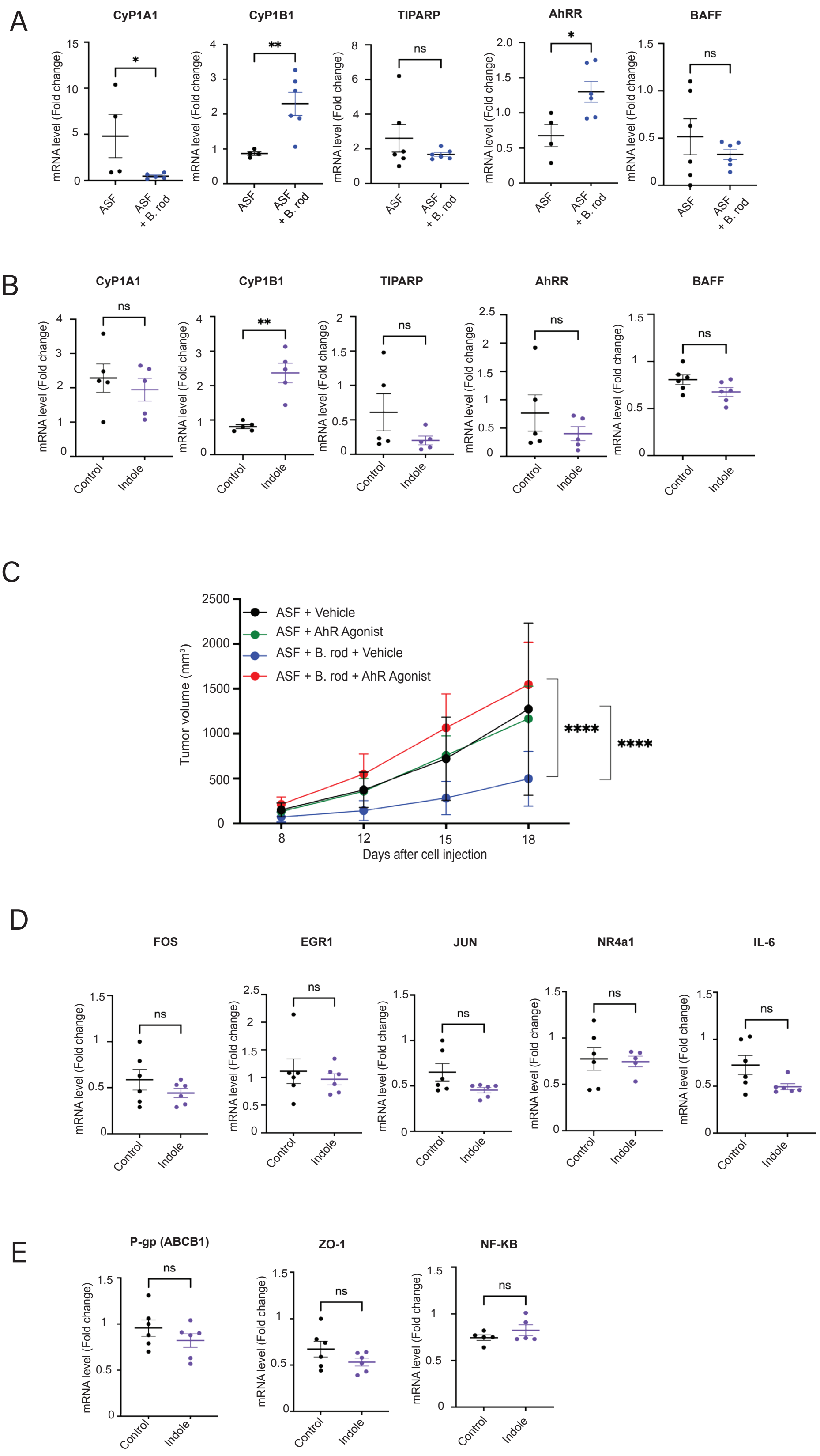


Diaz Olea et al., Sup Fig 6
